# Integrated spatial and single-cell transcriptomic analysis of aggressive glioblastoma growth dynamics

**DOI:** 10.64898/2026.05.11.724432

**Authors:** Clara Alves-Pereira, Gyeong Dae Kim, Ngima Sherpa, Kayla Colvin, Saad M. Khan, Khanh P. Phan, Anthony Z. Wang, Ian F. Dunn, Tanner Johanns, Erdyni Tsitsykov, Rupen Desai, Gavin P. Dunn, Allegra A. Petti

## Abstract

Glioblastoma (GBM) develops within a complex tumor ecosystem whose temporal dynamics remain poorly understood. Here, we performed longitudinal single-cell RNA sequencing and spatial transcriptomics across multiple timepoints in two widely used murine GBM models - CT2A and GL261 - which differ markedly in aggressiveness and response to immune checkpoint blockade. Tumor cell transcriptomes revealed model-specific programs: CT2A cells progressively upregulated epithelial-mesenchymal transition (EMT), non-classical MHC Class I, and progressively, hypoxia response pathways, resembling the human mesenchymal GBM cell state, while GL261 cells exhibited MHC Class II expression and developmental signatures resembling oligodendrocyte progenitor and astrocytic states. Ligand-receptor interaction analyses identified thrombospondins (*Thbs1*, *Thbs2*) and osteopontin (*Spp1*) as CT2A-specific tumor ligands mediating tumorigenic interactions with immune cells, with downstream targets enriched for EMT and TGF-β pathways. Conversely, the GL261 model presented a differential potential to engage neuronal and perivascular guidance networks, with Glutamate and L1 cell adhesion molecule (*L1cam*) as lead signaling partners. The CT2A immune compartment exhibited progressive microglia-to-macrophage phenotypic conversion, enhanced macrophage infiltration driven by *Spp1*, and elevated T cell exhaustion, while GL261 maintained a distinct adaptive immune communication hub via MHC class II-CD4 signaling. Elevated *THBS1*, *THBS2*, and *SPP1* expression correlated with poor survival in human GBM datasets. Together, these findings reveal divergent tumor-immune ecosystems in CT2A and GL261 that recapitulate distinct aspects of human GBM, with implications for therapeutic targeting.

## Introduction

Glioblastoma, the most common primary malignant brain tumor in adults, develops within a unique ecosystem characterized by immune dysfunction, hypoxia, necrosis, and microvascular proliferation. These features likely contribute to the failure of targeted treatments and immunotherapies that have been effective in other tumor types (Goto et al., 2023; Gschwandtner et al., 2019; Khan et al., 2024; X. Wu et al., 2024). However, the tumor microenvironment we observe at time of diagnosis upon resection is an evolved state representing the culmination of numerous sequential, reciprocal changes that have occurred throughout tumorigenesis. It is widely accepted that understanding the stepwise development of the tumor and its ecosystem can yield therapeutically important insights. In many cancer types, therefore, precursor lesions are actively studied, often at multiple time points during disease progression, to better understand these events (Chen and Lau, 2023). In GBM, however, no readily discernible premalignant lesion exists, precluding such studies in patients.

Syngeneic orthotopic rodent models of GBM can address this gap. Although inherently limited by their transplant origin rather than *de novo* tumorigenesis, they remain the most widely used preclinical GBM systems because they are immunocompetent, reproducible, and experimentally tractable. Among these, GL261 and CT2A are the most widely used. Both lines were derived through methylcholanthrene mutagenesis (Ausman et al., 1970; Zimmerman and Arnold, 1941) and yield 100% lethality when injected orthotopically into syngeneic C57Bl/6 mice. Both show high mutation burden relative to human GBM, but otherwise recapitulate hallmark GBM features including microvascular proliferation, necrosis, and infiltrative intracranial progression (Candolfi et al., 2007; Martínez-Murillo, 2007; Newcomb and Zagzag, 2009; Oh et al., 2014; Szatmári et al., 2006). However, the two differ markedly in natural history and immune phenotype. CT2A is more aggressive with a median survival 18-20 days compared to 25-30 days for GL261 (Haddad et al., 2021; Szatmári et al., 2006), and is generally refractory to immune checkpoint blockade (ICB), while GL261 frequently shows benefit (Genoud et al., 2018; Iorgulescu et al., 2023; Connor J Liu et al., 2020a; Reardon et al., 2016).

Here, we extend these studies by longitudinally performing single-cell and spatial transcriptomics (ST) in CT2A and GL261 to determine how the tumor microenvironment is remodeled in models that differ substantially in therapeutic responsiveness and immunologic state. Our data reveal disparate transcriptional programs, as well as the ligand-receptor interactions that may mediate the downstream effects of these programs on the tumor and immune microenvironment. These findings suggests that each tumor model captures complementary features of human GBMs.

## Results

### Characterization of tumor growth dynamics

Prior work has shown that CT2A tumors are more aggressive than GL261 tumors (Haddad et al., 2021; Connor J. Liu et al., 2020; Szatmári et al., 2006), suggesting differing growth dynamics of CT2A and GL261. To identify comparable time points for sampling, we utilized serial Magnetic Resonance Imaging (MRI) to generate growth curves for each model (Fig. 1B, Methods). However, results were inconsistent with disease developmental stage and timepoint (see Supplementary Note 1) and points of collection were therefore determined to represent developmental stage and terminal point. The following timepoints were selected for analysis: “Early” tumor establishment (day 7 for both models), “Middle” for early exponential growth (day 14 for GL261; day 10 for CT2A), and “Late” exponential growth (day 17 for GL261, day 15 for CT2A) (Fig. 1A).

**Figure 1.**
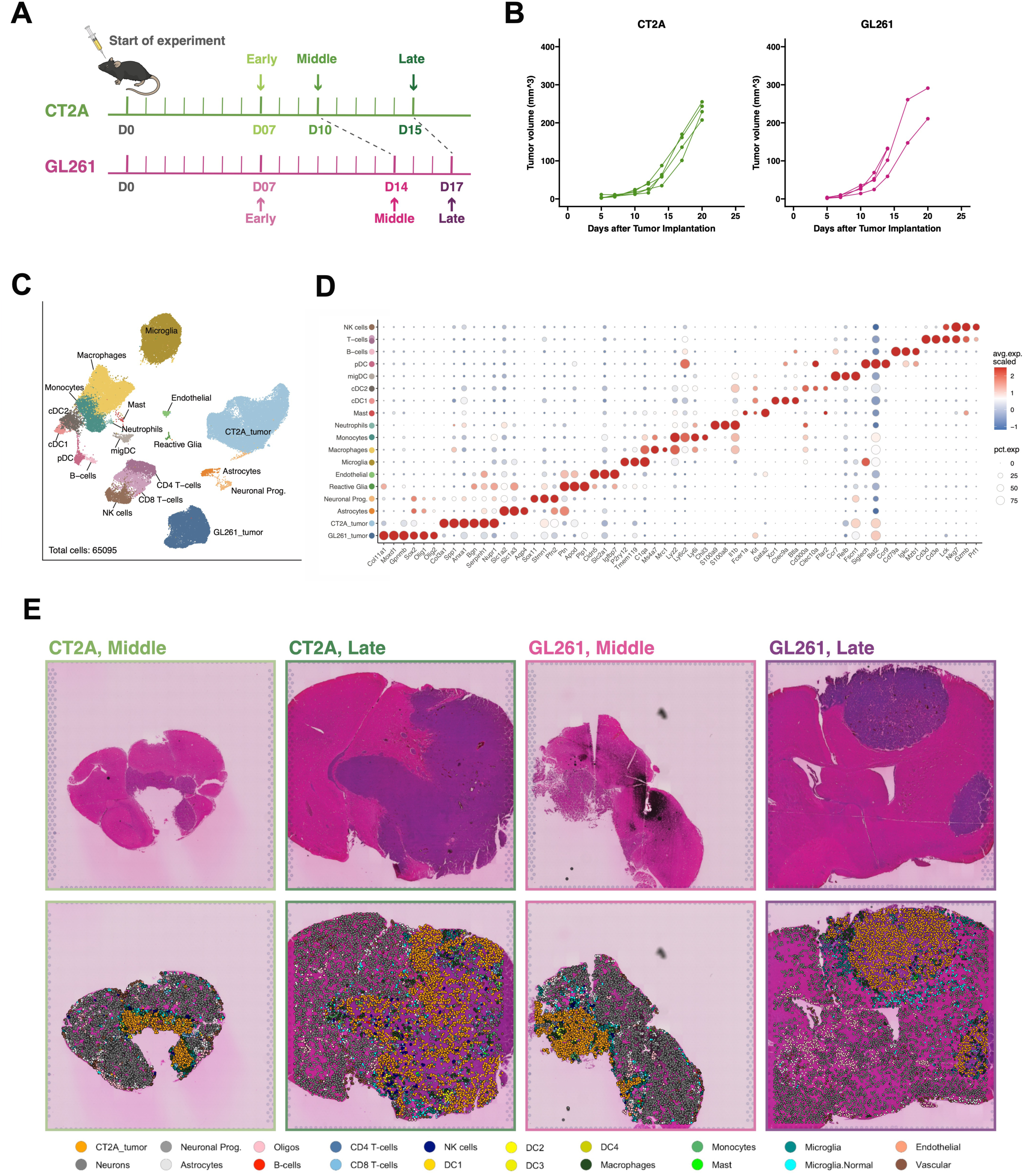
Sample preparation and profiling of scRNA-seq and spatial transcriptome datasets. **A.** Schematic of timecourses experiment experimental design. **B.** Tumor volumetric curves obtained from MRI of CT2A and GL261 mouse models over time. **C.** UMAP visualization of a total of 65,095 cells from CT2A and GL261 tumors developed from orthotopic injections and collected at 3 timepoints per model. For each timepoint, 5 tumors from 5 animals were pooled and independently processed for single-cell analysis (10X technology). **D.** Dot plot showing the scaled expression level of marker genes within each cluster. **E.** Hematoxylin and Eosin (H&E) staining of the four spatial samples (top) and cell type projection distribution (bottom) across time points. To build the scRNA-Seq reference for cell type deconvolution, a snRNA-Seq dataset of mouse normal brain was spiked-in onto the scRNA-Seq data. This merged reference was projected onto the spatial slides using the CellTrek package (Wei et al., 2022), allowing for the detection not only of tumor and immune cells, but also normal brain parenchyma cell populations.

### Temporal landscape of GBM progression using single-cell RNA-sequencing (scRNA-seq) and spatial transcriptomics (ST)

We performed scRNA-seq across at each timepoint, pooling resected tumor tissue from 5 animals per timepoint. We also performed ST at the middle and late timepoints, for both tumor models (Methods). After quality control, dimensionality reduction, and marker-based cell typing of the merged scRNA-seq data set (Methods), we identified 18 broad cell types, including two populations of tumor cells (representing CT2A and GL261), astrocytes, neuronal progenitors, microglia, reactive glial cells, endothelial cells, and diverse immune cells (Fig. 1C,D). We also examined the overall distribution of cell types across different timepoints (Fig S1A,B). Non-tumor and non-immune populations (e.g. reactive glia and endothelial cells) were primarily detected in low numbers at the earliest stage of the disease, disappearing as the tumor progressed and expanded to become the predominant cellular component. Likewise, in the immune compartment, the brain-native microglia population is drastically reduced along tumor progression, but partially replaced by infiltrating macrophage populations. Interestingly, monocyte proportions are also greater at the earliest timepoint (Fig. S1A,B). Visium ST data was filtered for high-quality spots and deconvolved using CellTrek (See Supplementary Note 2). The ST data revealed regions of dense tumor cells corresponding to visible tumor in the H&E images, as well as spatially segregated populations of diverse nonmalignant cells (Fig. 1E). Immune cells displayed time-dependent changes in spatial distribution, with greater tumor infiltration at the late time point, suggesting dynamic interactions between tumor and immune cells (Fig. 1E, Fig. S3C,F, Fig. S4C,F).

### CT2A and GL261 models recapitulate divergent mesenchymal and pro-neural human glioblastoma states

The CT2A and GL261 murine models represent two distinct poles of glioblastoma biology: work from our group and others have shown that GL261 is partially responsive to immune checkpoint inhibition (ICI), while CT2A is highly aggressive and largely refractory to immunotherapy (Genoud et al., 2018; Haddad et al., 2021; Iorgulescu et al., 2023; Connor J Liu et al., 2020b; Reardon et al., 2016; Szatmári et al., 2006). To identify transcriptional signatures underlying these differences at the untreated baseline, we first compared CT2A and GL261 tumor cells using standard tests of differential gene expression (Fig. 2A, Table S1, Methods). This revealed striking differences: genes in pathways associated with tumor aggressiveness (e.g. epithelial mesenchymal transition (EMT), angiogenesis, morphogenesis (Hox genes), pro-invasive programs (Hippo-YAP/TAZ), hypoxia response, inflammatory programs related to wound-healing) were more highly expressed in CT2A, whereas genes involved in neuroglial development, myelination and antigen presentation were more highly expressed in GL261 (Fig. 2A, Fig. S2A, Supplementary Table 1). We scored the tumor cells for the enriched pathways in the two models, redemonstrating these results with an expected temporal evolution (Fig S2B). These distinct but rich transcriptional signatures suggest that the two mouse models can recapitulate the cellular heterogeneity observed in human patients (Kirishima et al., 2024; Parker et al., 2015; Patel et al., 2014). To test this, we mapped our data onto established human glioblastoma meta-modules. We found that CT2A cells align closely with the Mesenchymal (MES) state, a phenotype historically associated with high grade, poor prognosis, and a robust inflammatory niche. In contrast, GL261 cells resemble the Oligodendrocyte Progenitor (OPC) and Astrocytic (AC) states, which are characterized by neuro-developmental programs and a more permissive immune environment (Fig. 2B). Relative metaprogram scores plotted over time also revealed that CT2A cells strictly score MES1+MES2 (MES) with no variation over time - revealing a much higher strength of this meta-module in CT2A cells, when measured against the other metaprograms - while in GL261 tumors the metaprogram signatures have a more hybrid composition, scoring MES, AC and OPC, with OPC scores increasing over time (Fig 2B, bottom panel).

**Figure 2.**
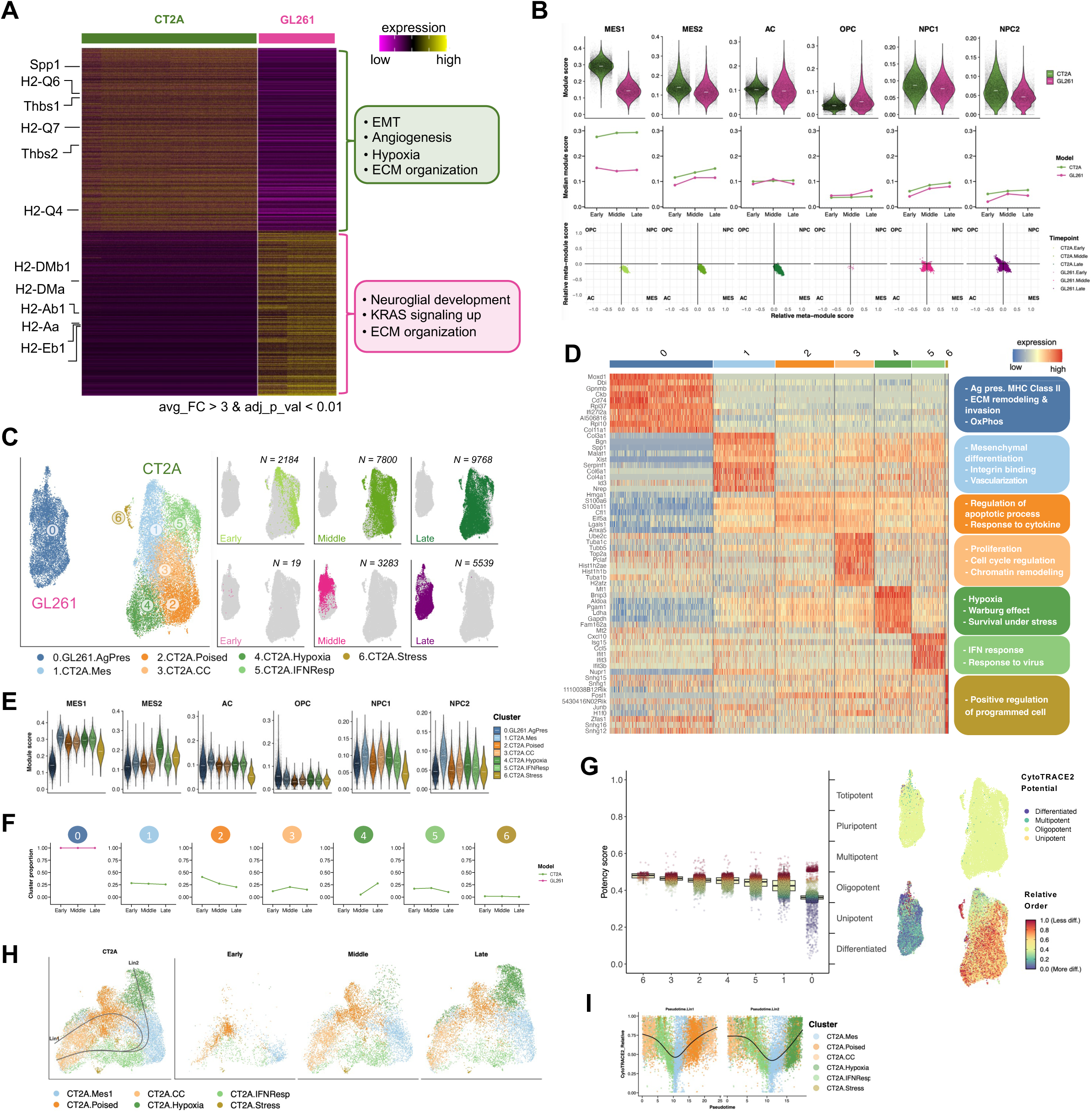
Transcriptional heterogeneity and human cell-state modeling of CT2A and GL261 tumor cells. **A.** Differential gene expression (DGE) and pathway enrichment between CT2A and GL261 tumor cells. **B.** Transcriptional scoring of tumor cells based on human GBM meta-modules (Neftel et al., 2019). Violin plots (top) illustrate model-specific scores per meta-module; line plots (middle) represent temporal evolution of scores; quadrant plots (bottom) illustrate relative distributions across four merged signatures: OPC, NPC (NPC1+NPC2), MES (MES1+MES2), and AC. **C.** UMAP projections of tumor cell sub-clusters. Individual UMAPs are colored by model per timepoint. **D.** Heatmap transcriptional profiles and functional enrichment of tumor cell sub-clusters. Only the top 10 DE genes shown (left) but overrepresentation analysis was performed with the 100 most differentially expressed genes in each cluster when compared to all other clusters. **E.** Violin plots showing the Neftel scores across tumor clusters. **F.** Temporal dynamics of tumor cluster proportions across sampled timepoints. **G.** Differentiation potential scoring of tumor clusters via CytoTRACE2. Boxplot showing stemness scores per tumor cluster (left) and UMAP projections (right) represent absolute and relative differentiation states. **H.** Pseudotemporal trajectory reconstruction of CT2A tumor cells. Principal curves (Lineage 1 and Lineage 2) were calculated via Slingshot; sub-panels illustrate trajectories across discrete timepoints. **I.** Dynamic CytoTRACE2 scores along reconstructed pseudotime for CT2A tumor lineages.

To resolve how these transcriptional states manifest as intratumoral heterogeneity, we sub-clustered tumor cells from both models (Fig. 2C, Methods). This analysis yielded seven populations exhibiting distinct transcriptional programs and biological pathways (Fig. 2C,D). These confirmed the coarse distinctions between CT2A and GL261 identified in Fig. 2A, and, surprisingly, revealed notable intratumoral heterogeneity within CT2A. Specifically, GL261 resolved into a single cluster (C0) enriched for glial cell differentiation genes, ECM organization, and MHC Class II antigen presentation. In alternative iterations, GL261 resolved into a maximum of 3 clusters, however, DE analysis suggested no major structure between these clusters at the gene expression level leading to segregations of major functional states. On the other hand, CT2A was composed of functionally diverse clusters enriched for proliferative states (C3), interferon I/II response (C5), EMT (C1), hypoxia response (C4) and stress (C6). Notably, the hypoxia-related cluster (C4) in CT2A progressively expanded over time (Fig. 2F). Comparing the median scores of Hallmark signatures in the clustered dataset revealed patterns hidden in the bulk analysis. For example, while IFNγ scores were overall lower in CT2A and decrease over time in both models (Fig. S2 B), the score is consistently elevated over time in the IFN-responsive CT2A cluster 5 (Fig. S2 C).

To further identify integrated biological programs, we used hdWGCNA (Morabito et al., 2023) which identifies modules of co-expressed genes. CT2A tumors showed upregulation of modules enriched for mTOR, HIF1A, and Wnt/beta-catenin signaling pathways, consistent with transcriptional programs associated with aggressiveness and reduced cell differentiation (Fig. S2D). To independently assess differentiation state, we performed CytoTRACE2 analysis (see Supplementary Note 2). This analysis showed higher potency scores on average for CT2A than GL261 cells, consistent with an overall less differentiated transcriptional state overall (Fig. 2G). These results align with the classification of CT2A as a mesenchymal model and GL261 as a more differentiated, pro-neural model. However, GL261 included a small subset of cells with higher inferred potency (Fig. 2G) potentially consistent with previously described stem-like or tumor-propagating subpopulations (Wu et al., 2008). Across both models, inferred potency scores changed only modestly with tumor progression (Fig. S2E), supporting relative stability of differentiation states.

To resolve the dynamic transitions between functional states, we performed pseudo-temporal trajectory analysis on CT2A tumor cells. A branched developmental architecture was inferred, originating from a proliferative state that bifurcated at the transition between the interferon response cluster and the mesenchymal clusters (Fig. 2H). Lineage 1 progressed toward a proliferative cell state via a transitional ’Poised’ cluster, which exhibited low expression of specialized markers (Fig 2D). In contrast, Lineage 2 followed a trajectory suggestive of reactive adaptation, passing through IFN-response and Mesenchymal clusters before culminating in a terminal Hypoxia-responsive state (Fig. 2H). Curiously, CytoTRACE2 scores remained high at both ends of these lineages - the proliferative and hypoxic clusters - while dipping in the mesenchymal intermediate (Fig. 2I). The results suggest a model of niche-driven dedifferentiation, where the extreme stress of the hypoxic environment reinforces a stem-like state in CT2A cells, while, counter-intuitively, the “mesenchymal” transition cluster represents a more specialized phenotype. This branched trajectory highlights the plasticity of CT2A, possibly allowing it to simultaneously pursue rapid expansion (lineage 1) and stress-adapted survival (lineage 2).

To map the tumor states within the intact tumor architecture we used Visium spatial transcriptomics (Fig. S3, S4). Across two timepoints (Middle, Late), spots were deconvolved into their constituent cells using the CellTrek package (Methods, Supplemental Note 2) (Fig. 1E). Tumor cells and healthy neurons mapped as expected in the tumor and the parenchyma regions defined by H&E (Fig. 1E), while immune cells were mostly located in the interfaces between these regions. Cell deconvolution combined with spot clustering analysis revealed that clusters with tumor-enriched gene signatures were predominantly enriched for tumor cells and, similarly, clusters mostly enriched for neuronal and glial gene signatures were enriched for neuronal and glial deconvolved cell types. Those clusters were predominant, while only a few clusters were dominated by immune and vascular cell types (Fig. S3, S4). In late-stage CT2A, tumor-enriched clusters exhibited strong EMT- and hypoxia-associated signals (Fig. S3E). In contrast, tumor clusters in GL261 were primarily characterized by cell cycle-related signals (Fig. S4E). Notably, unlike CT2A middle or either GL261 samples, CT2A late samples showed co-localization of tumor cells with neurons and multiple immune cell types, including macrophages and T cells, within a single cluster (Fig. S3C,F, S4C,F). This observation hints at the infiltrative nature of the CT2A sample, especially at later stages.

### Thrombospondin and osteopontin are key ligands in CT2A-specific interactions and correlate with EMT and hypoxia

To prioritize model-specific L–R interactions, we corroborated the CellChat results using an independent ligand–receptor inference algorithm, CellPhoneDB (Troulé et al., 2025). We intersected these results with top DEGs and identified thrombospondins (*Thbs1*, *Thbs2*), osteopontin (*Spp1*) and the collagens (Col6a1, Col4a1) as the dominant ligands mediating CT2A tumor-specific communications with relatively higher interaction strength for Spp1 and Thbs1/2 (Fig. 3A,B,C; Fig. S6B,C). ST confirmed the enrichment of these transcripts in CT2A (Fig. 3D). Spatial L-R analysis using Niches (Raredon et al., 2023) (Methods) show that the interactions of Thbs1 and Thbs2 with Cd47, Sdc1 and Sdc4 were stronger in CT2A than in GL261, and stable across timepoints (Fig. 3D).

**Figure 3.**
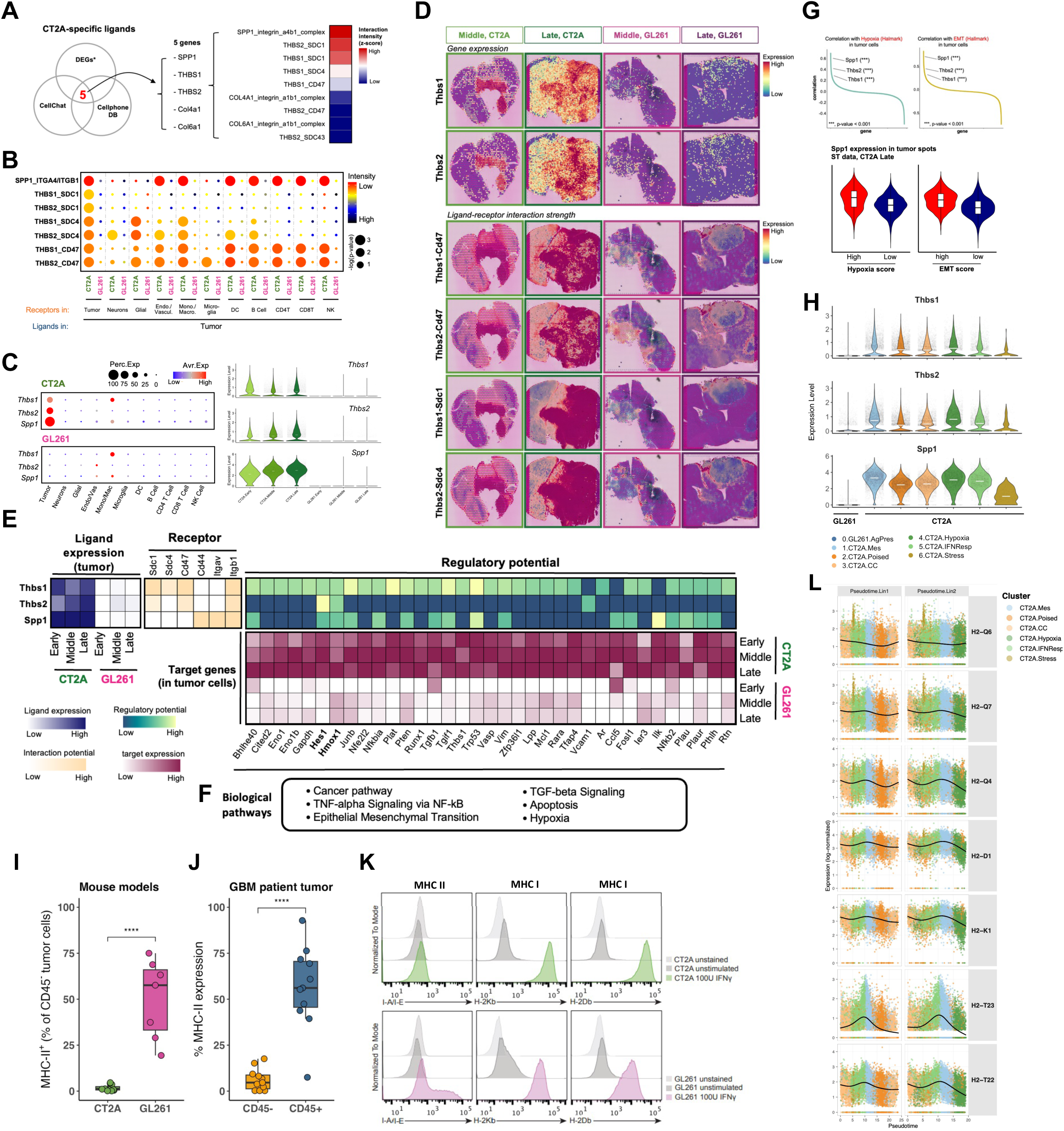
Divergent ligand-receptor interactions and mechanisms of immune evasion in CT2A and GL261. **A.** Venn diagram (left) identifying CT2A-enriched ligands consistently detected by CellChat and CellPhoneDB; heatmap (right) indicating interaction intensities for these ligands. **B.** Interaction probabilities for tumor cell secreted ligands across all recipient cell types; dot size and color represent -log_10_(p-value) and interaction intensity, respectively. **C.** Expression of *Thbs1, Thbs2* and *Spp1* across scRNA-seq cell types (left) and temporal expression profiles in tumor cells (right). **D.** Spatial transcriptomic feature plots illustrating *Thbs1/2* expression (top) and corresponding ligand-receptor interaction strength (bottom) in CT2A and GL261 across timepoints. **E.** Integrated analysis of *Thbs1*, *Thbs2*, and *Spp1* autocrine (tumor -> tumor) regulatory potential. Heatmaps display temporal ligand expression in tumor cells (top left), predicted regulatory potential for downstream targets (top right middle), and target gene expression across time points (bottom right). **F.** Pathway enrichment analysis of the target genes identified in (E). **G.** Correlation between hypoxia and EMT biological scores and the tumor cell transcriptome from the single cell dataset. **H.** Expression of *Thbs1*, *Thbs2*, and *Spp1* across identified tumor sub-clusters. **I.** Flow cytometric analysis of MHC-II (I-A/I-E) expression in the CD45-negative fraction of CT2A and GL261 tumors. **J.** MHC-II expression in the CD45-negative fraction of resected human GBM tumors (*n=11*). For (I) and (J), significance was determined by Wilcoxon rank-sum test (R base). **K.** Normalized expression of classical and non-classical MHC-I genes along CT2A tumor cell pseudotime as in Fig. 2I.

We next used NicheNet (Browaeys et al., 2020) to investigate potential downstream target genes regulated by thrombospondin and osteopontin interactions. Given their dominant enrichment in CT2A, we reasoned these ligands may partially drive the gene expression programs that distinguish CT2A from GL261. The analysis predicted that *Thbs1/2* and *Spp1* likely regulate a gene sets related to EMT, TGF-β, and hypoxia pathways (Fig. 3E,F). Notably, two genes, *Hmox1* and *Hes1*, emerged as common downstream targets of all three ligand/receptor regulons (Fig. 3E). Both genes have documented roles in tumor progression in response to hypoxia, but act through different mechanisms: *Hes1* promotes glioma stem cell proliferation and EMT (Qiang et al., 2012), while *Hmox1* (encoding HO-1) suppresses anti-tumor immunity (Lee et al., 1997). The influence of *Hmox1* on the tumor immune microenvironment is multifaceted. Specifically, HO-1 has been shown to promote M2 polarization, enhance TAM infiltration, and suppress anti-tumor cytokine production by T lymphocytes (Alaluf et al., 2020; Luu Hoang et al., 2021). Importantly, HO-1 expression negatively correlates with MHC class II expression in the CNS, suggesting a mechanistic link between hypoxia and immune recognition (Alaluf et al., 2020; Chora et al., 2007). Our hdWGCNA analysis further supported a link with response to hypoxia, placing *Thbs2* within the hypoxia-associated module and *Spp1* within the EMT-associated module (Fig. S2D). All three ligands (*Thbs1*, *Thbs2*, *Spp1*) correlated positively with hypoxia and EMT scores (Fig. 3G) and were enriched in the mesenchymal and hypoxic tumor clusters (Fig 3H).

Finally, we confirmed a correlation between survival and expression of *Thbs1*, *Thbs2*, and *Spp1* in human transcriptomic and proteomic datasets obtained from The Human Protein Atlas (Uhlen et al., 2017; Uhlén et al., 2015). This analysis revealed that elevated expression (above the median values) of these genes in human GBM is associated with a poorer survival (Fig. S6D), suggesting that they may be contributing to the more aggressive phenotype of CT2A. Taken together, these findings indicate that CT2A tumor cells, or a subpopulation thereof, are hypoxic and EMT-like, and participate in pro-tumorigenic intercellular interactions with the TME mediated by *Thbs1*, *Thbs2*, and *Spp1*.

### Divergent antigen presentation programs in CT2A and GL261 models

Our transcriptional profiling highlighted a stark divergence in immune signaling pathways between the two models (Fig. 2A). Gene Set Enrichment Analysis (GSEA) revealed that while CT2A was enriched for TGF-β signaling, TNF via NF-κB, and MHC-I pathways, GL261 was characterized by MHC-II signaling, IFNγ response, and allograft rejection (Fig. S2B). In CT2A, the upregulation of TGF-β pathway coincided with upregulation of TNFα-driven inflammatory signaling (Fig. S2B,C) and increased *Ccl2* expression, a potent chemoattractant for myeloid cells, which contribute to an immunosuppressive environment. GL261 cells expressed MHC-II molecules (Fig. S7B) and remarkably, in our L-R analysis prioritization strategy, MHC-II genes and collagen genes were dominant interactions in GL261 (Fig. S7A,C,D). MHC-II is rarely expressed in tumor cells, and is usually restricted to antigen presenting cells (APCs). However, *MHC-II* expression is typically associated with cancers where ICI is historically effective (Alspach et al., 2019; Christopher et al., 2018; Johnson et al., 2020). We also observed *Irf8,* a master transcription factor typically restricted to the myeloid lineage, expression in GL261 tumor cells (Fig. S7B). Recent evidence has shown that *Irf8* expression in immunocompetent GBM models can be a byproduct of immune pressure triggering tumor cells to adopt an immunosuppressive myeloid-like transcriptional program (Gangoso et al., 2021). MHC-II expression in GL261 could be supported by *Ciita* (Class II Major Histocompatibility Complex Transactivator) jointly with *Irf8*. Both transcription factor genes are expressed at low levels in GL261, but not in CT2A (Fig. S7B).

To orthogonally validate the MHC-II expression in GL261 tumor cells at the protein level, we performed flow cytometry on brain tumor samples from both models. From *ex viv*o tumors, the percentage of MHC-II expressing CD45-negative cells was significantly greater in GL261 tumors compared to CT2A tumors (Fig. 3I). We extended this analysis to freshly resected patient GBM samples. While most tumors exhibited little to no expression, some tumors had some expression of MHC-II, highlighting that MHC-II can be expressed in patient tumors (Fig. 3J). It is thus worth exploring how MHC-II expression on tumors predicts survival. To determine if lack of MHC-II expression in CT2A was a plastic response or a fixed trait, we stimulated both lines with IFNγ *in vitro*. While a subset of GL261 cells exhibited expression of MHC-II upon stimulation of IFN-g, CT2A did not show induction of MHC-II (Fig. 3K), suggesting a cell-intrinsic barrier to MHC-II presentation. However, both cell lines expressed MHC-I (H2-Kb and H2-Db) upon IFN-g stimulation, showing that the MHC-I protein expression at the cell surface in CT2A can be fully rescued.

The upregulation of MHC-I pathways in CT2A was is notable, because defects in the class I antigen presenting machinery, affecting *Tap1* and *Psmb8* genes are well described for CT2A cells (Iorgulescu et al., 2023). In our scRNA-Seq dataset, when expressed in CT2A, MHC-I expression is relatively higher than in GL261, including the *B2m* gene (Fig. S7B). Using pseudotime trajectory analysis in CT2A tumor cells, we observed that both classical (H2-K1, H2-D1) and non-classical (H2-T23, H2-T22, H2-Q) MHC-I genes frequently exhibit a peak expression along the mesenchymal ordered cells and a noticeable decline in expression, in Lineage 2, as tumor cells transition into the hypoxic cluster (Fig. 3L).

### Progressive macrophage expansion, microglial reprogramming, and hypoxia define the aggressive CT2A myeloid landscape

Given the strong immunomodulatory signature of CT2A tumor cells, we next characterized the immune cell compartments in both murine models. Sub-clustering of the immune compartment yielded 11 clusters, with myeloid cells showing the greatest shifts in abundance (Fig. 4A, B). Macrophage proportions progressively expanded while microglia declined over time, with more striking changes in CT2A (Fig. 4B). At the early timepoint, CT2A already shows higher macrophage proportion, while GL261 is dominated by tissue-resident microglia. Interestingly, monocyte proportions decreased over time in GL261 but remained stable in CT2A, consistent with sustained recruitment and rapid differentiation into macrophages in the more aggressive model (Fig. 4B).

**Figure 4.**
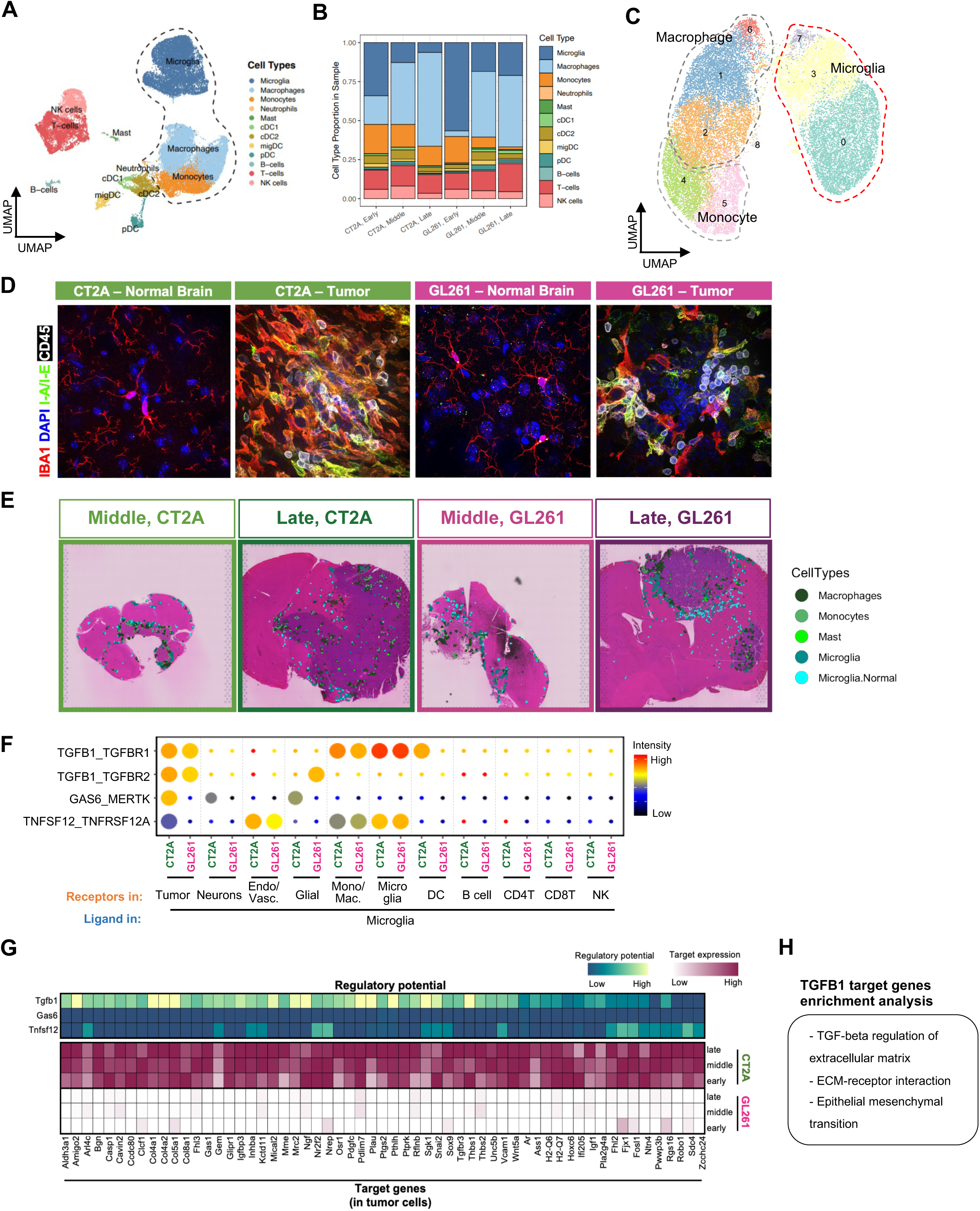
Spatiotemporal remodeling of myeloid cells and their interactions in CT2A and GL261. **A.** UMAP projection of the immune cell compartment. **B.** Proportional changes in immune cell populations across sampled timepoints. **C.** Sub-clustering of the myeloid compartment. UMAP including macrophages, monocytes, and microglia. **D.** Representative immunofluorescence (IF) micrographs of late-stage orthotopic CT2A (left) and GL261 (right) tumors. Myeloid cells are stained with IBA1 (red), alongside I-A/I-E (MHC-II, green), CD45 (white), and DAPI nuclei counterstain (blue); magnification, 100X. **E.** Spatial mapping of myeloid populations. Distribution of myeloid sub-types in middle and late-stage tumors following spatial deconvolution. **F.** Microglia-to-microenvironment ligand-receptor interactions. Dot size and color represent -log(p-value) and interaction intensity, respectively, between microglial ligands and recipient cell receptors. **G.** Predicted regulatory potential of microglial-derived ligands. Heatmaps illustrate the likelihood of Tgfb1, Gas6, and Tnfsf12 regulating specific downstream target genes (top) and the temporal expression of those targets in tumor cells (bottom). **H.** Pathway enrichment of microglial-regulated tumor cell targets identified in (G).

Macrophage and microglia phenotypes are notoriously challenging to distinguish in gliomas, but scRNA-seq provides complementary molecular resolution despite limiting accurate cell quantification. To compare myeloid infiltration histologically, we performed immunofluorescence (IF) at the late timepoint in CT2A and GL261 using IBA (pan-myeloid) and MHC-II (enriched in macrophages) (Fig. 4D). CT2A showed higher abundance of IBA+ cells than GL261 (Fig. 4D). Morphologically, IBA+ cells in CT2A adopted an amoeboid, activated-like state compared to the ramified morphology typical of homeostatic microglia (as seen in the unaffected contralateral hemisphere), a shift less pronounced in GL261 (in Fig. 4D). These observations suggest a stronger reprogramming of microglia, higher infiltration of macrophages, or both, in CT2A. To test this, we subsetted myeloid cells from the scRNA-seq data. Batch-corrected UMAP showed a clear separation of microglia and monocyte/macrophage clusters (Fig. 4C, S8A,B). We then scored the microglia meta-cluster with the macrophage signature and did not detect differences between models (Fig. S8E). However, microglia homeostatic markers (*P2ry12* and *Tmem119*) decreased over time, and macrophage/activation markers (*Lyz2* and *Cd14*) increased (Fig. S8F). At the sub-cluster level (microglia clusters C0,C3,C7) C3 showed higher macrophage scores (Fig. S8G), lower microglia homeostatic markers and higher macrophage/activation markers (Fig. S8H). Notably, C3 expanded over time, particularly in CT2A, where it nearly dominated the microglial compartment at the late stage (Fig. S8I). These observations are consistent with a model where gliomas drive a progressive microglial reprogramming, more pronounced in the aggressive CT2A model, while GL261 retains a more homeostatic microglial state.

Among interactions that may have contributed to immunosuppressive environment, the probability of TNFSF12 interactions, which are known to drive tumor progression and immune evasion (Chen, 2024; Culp et al., 2010), increased in both models at later time points when sent by microglia and received by other myeloid cells, and *TGFB1* when interacting with tumor (Fig. 4F, S9J). In CT2A tumor cells, *Tgfb1* sent by microglia likely contributed to the mesenchymal phenotype (Fig. 4G,H). Within the immune compartment, microglia interacted not only with macrophages, but also with other immune cells including CD4 T cells, CD8 T cells and NK cells through inhibitory interactions such as *Lgals9*:*Havcr2*, *Cd274*:*Pdcd1*, and *Nectin2*:*Tigit* (Fig. S9J), likely contributing to immune suppression within the tumor microenvironment (Chauvin and Zarour, 2020; He and Xu, 2020; Jin et al., 2023; Kandel et al., 2021; Wienke et al., 2024).

Like microglia, macrophages in CT2A exhibited potentially immunosuppressive interactions with tumor cells and immune cells (Fig. 5B). Interactions mediated by *Ccl7*, *Cxcl14*, and *Ccl2* were connecting other immune cells, significantly more abundant in CT2A, and increased over time (Fig. 5B and S10A). Given their contribution to macrophage infiltration and transition into tumor-associated macrophages (TAMs), these CT2A-specific myeloid-mediated interactions are likely to further accelerate tumor progression (Goto et al., 2023; Gschwandtner et al., 2019; Khan et al., 2024; X. Wu et al., 2024). In addition, Tgfb1-Tgfbr1, Lgals9-Havcr2 and Cd274-Pd1 interactions, which are known to contribute to a pro-tumorigenic microenvironment, were predicted to be relatively stronger in CT2A (Fig. 5B and S10A,B). TGF-β-mediated interactions between macrophages and tumor cells were more prominent in CT2A, especially at early timepoints, as confirmed by both scRNA-seq and spatial data (Fig. 5C-last row and S10A), further contributing to the tumor phenotype (Fig. 4G). Additionally, macrophages in CT2A demonstrated stronger *Lgals9*:*Havcr2* and *Cd274*:*Pdcd1* interactions (Fig. 5B and S10A), which have been implicated in immune suppression and M2 polarization (He and Xu, 2020; Jin et al., 2023; Kandel et al., 2021).

**Figure 5.**
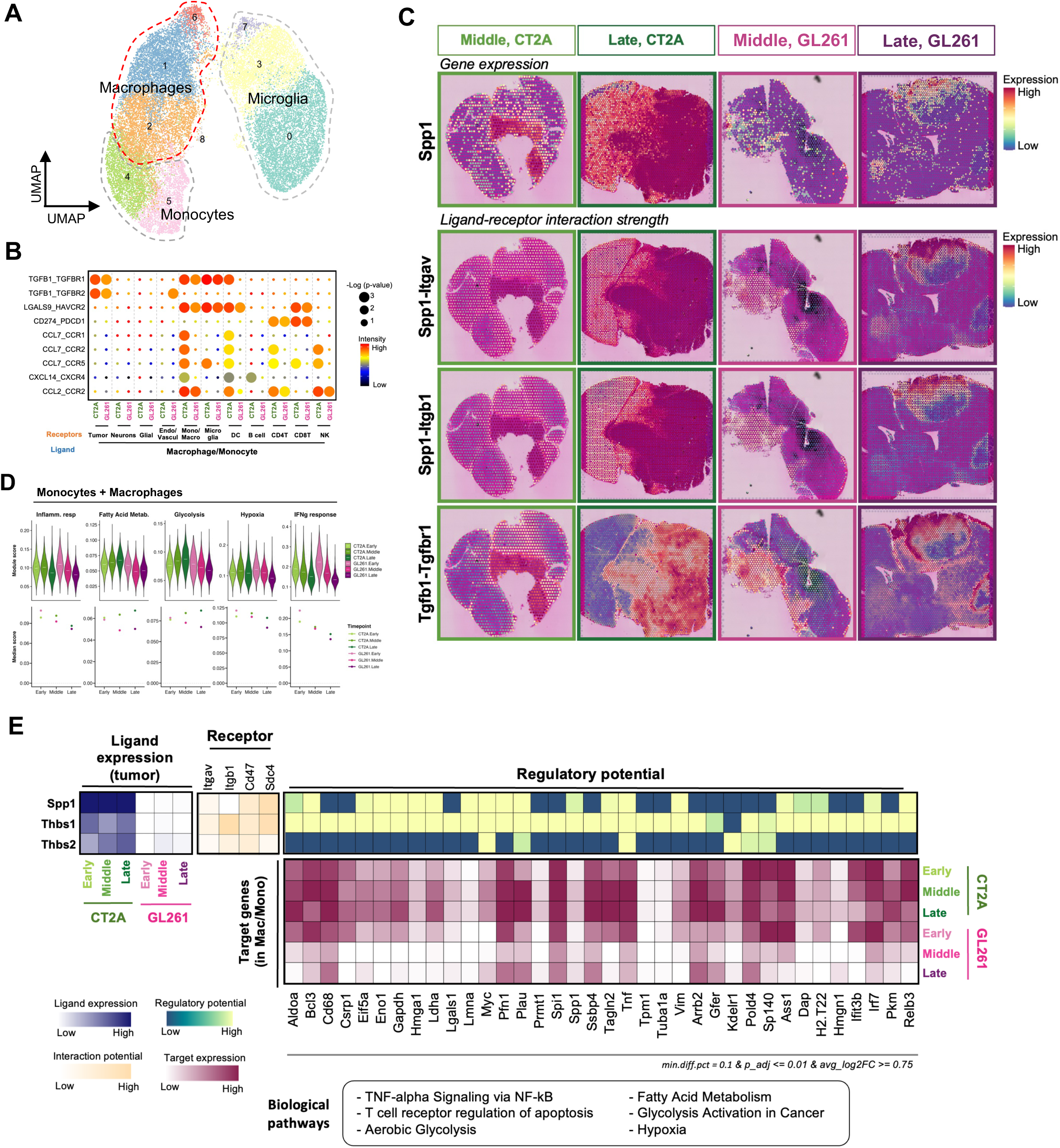
Spatiotemporal dynamics and transcriptional reprogramming of tumor-associated macrophages. **A.** Sub-clustering and identification of myeloid populations. the macrophage cluster is delineated by a dashed red boundary. **B.** Spatiotemporal macrophage-to-microenvironment ligand-receptor interactions. Dot size and color represent –log_10_(p-value) and interaction intensity, respectively, for macrophage-derived ligands across sampled timepoints. **C.** Spatial mapping of *Spp1* and *Tgfb1* signaling. Feature plots illustrate the spatial distribution of *Spp1* expression (top) and predicted interaction strengths for *Spp1-Itgav, Spp1-Itgb1,* and *Tgfb1-Tgfbr1* (bottom) in CT2A and GL261 tumors. **D.** Temporal variation in MSigDB Hallmark pathway scores. Signatures were calculated for the combined monocyte and macrophage compartment across tumor progression. **E.** Integrated analysis of tumor-to-macrophage signaling. Heatmaps illustrate the predicted regulatory potential of tumor-derived ligands (*Thbs1, Thbs2, Spp1*) and the corresponding temporal expression of predicted downstream target genes within the macrophage population.

In summary, the CT2A myeloid landscape is defined by an early and aggressive recruitment of monocyte-derived macrophages and a corresponding loss of microglial homeostatic identity. Driven by tumor-derived signals such as *Spp1* and *Thbs1/2* (Fig. 3B), these populations likely converge into an immunosuppressive niche that facilitates tumor progression and is associated with reduced T-cell infiltration. To characterize the effect of this inferred immunosuppressive state in the CT2A myeloid cells, we performed differential gene expression comparing a meta-cluster of macrophages and monocytes (we will transiently call this subset Monocyte-derived Tumor Associated Macrophages, MoTAMs) between the two models. Intriguingly, this analysis did not immediately highlight myeloid signatures of anti-tumor inflammation *vs.* pro-tumor immunosuppression (historically defined by a M1 vs. M2 states). Tumor associated macrophages are known to exhibit a complex multitude of immunomodulatory states beyond M1/M2 (Bied et al., 2023; Christofides et al., 2022). We have thus scored the CT2A and GL261 MoTAMs subset with the myeloid meta-programs defined for human GBM-associated macrophages by Miller *et al* (Miller et al., 2025). The results showed that, surprisingly, apart from a moderate higher inflammatory profile at earlier stages in GL261 MoTAMs, CT2A and GL261 immunomodulatory phenotypes are remarkably similar and even share patterns of progression over time (increased immunosuppression, transient increase of systemic inflammatory module, and later stages of decrease, Fig. S10C). This analysis highlights the presence of a common dysfunctional immunomodulatory myeloid compartment in both models. The DE and scoring analysis revealed, however, differences in metabolic and hypoxia programs (Fig. 5D). Interestingly, these differences evolved in diverging directions: while CT2A myeloid cells saw their hypoxic levels increase over time, GL261 stared with equivalent scores at early timepoints, which decreased over time (Fig. 5D). NicheNet analysis confirmed that *Spp1*, *Thbs1* and *Thbs2* have a positive regulatory potential for targets enriched in glycolytic and fatty acid metabolism as well as hypoxia in CT2A MoTAMs (Fig. 5E). While *Spp1* and *Thbs1* shared regulatory potential for glycolytic pathways, these three ligands share a common regulatory focus on key tumor-promoting targets - including *Plau*, *Pfn1*, and *Tnf* - all of which were expressed at higher levels in CT2A and increased as the tumor progressed. These three markers start to reveal how, in CT2A, tumor-educated macrophages differentially participate in enhanced pro-tumor functions relative to GL261: by upregulating *Plau* and *Pfn1,* macrophages promote both ECM proteolysis (Fleetwood et al., 2014), impact cell motility including, potentially, further macrophage infiltration and T cell exclusion (Gau et al., 2023; Schoppmeyer et al., 2017; Yun et al., 2011) and angiogenesis (Fan et al., 2014). Additionally, the temporal increase in *Tnf* expression indicates a state of chronic inflammation that may promote endothelial activation, aberrant angiogenesis, and enhanced tumor migration (Li et al., 2017)(Wei et al., 2021). Taken together, these results suggest a model where the *Thbs*/*Spp1* signaling axis in CT2A locks the myeloid compartment into a progressive hypoxic, glycolytic and invasive state. Conversely, the GL261 model appears capable of resolving its initial hypoxic environment (Fig. 5D). This metabolic resolution, combined with the sustained antigen presentation capabilities we observed previously, may provide a vulnerability that is exploited by ICI treatment in the GL261 model.

### Lymphocyte heterogeneity and immune dysfunction in CT2A

To investigate whether the aggressive CT2A niche impacts adaptive immunity, we analyzed the lymphocyte compartment in isolation and identified nine clusters (Fig. 6A). Each cluster was annotated based on representative marker genes (Fig. S11A). While total lymphocyte proportions were similar, CD4+ T cells were consistently more abundant in GL261 at the early timepoint (Fig. 6B). This observation aligns with our previous finding that CD4+ T cells are essential for ICI response in the GL261 model (Khan et al., 2022). In both models, infiltrating CD4 and CD8 T expressed canonical exhaustion markers, including *Pdcd1*, *Lag3*, and *Ctla4* (Fig. 6C and S11B). Notably, *Pdcd1*, a key regulator of immune checkpoint inhibitors, was expressed in both CD8 and CD4 T cells (Fig. 6C). However, the intensity of inhibitory signaling was markedly higher in the CT2A model. Specifically, PD-1/PD-L1 (Pdcd1/Cd274) receptor-ligand interactions were relatively higher in CT2A, particularly in CD8 T cells, and progressively increased over time irrespective of sender cell type (Fig. 6D and S11C, D). This suggests that while both tumors are "exhausted," the CT2A microenvironment imposes a more dense and consistent inhibitory checkpoint landscape.

**Figure 6.**
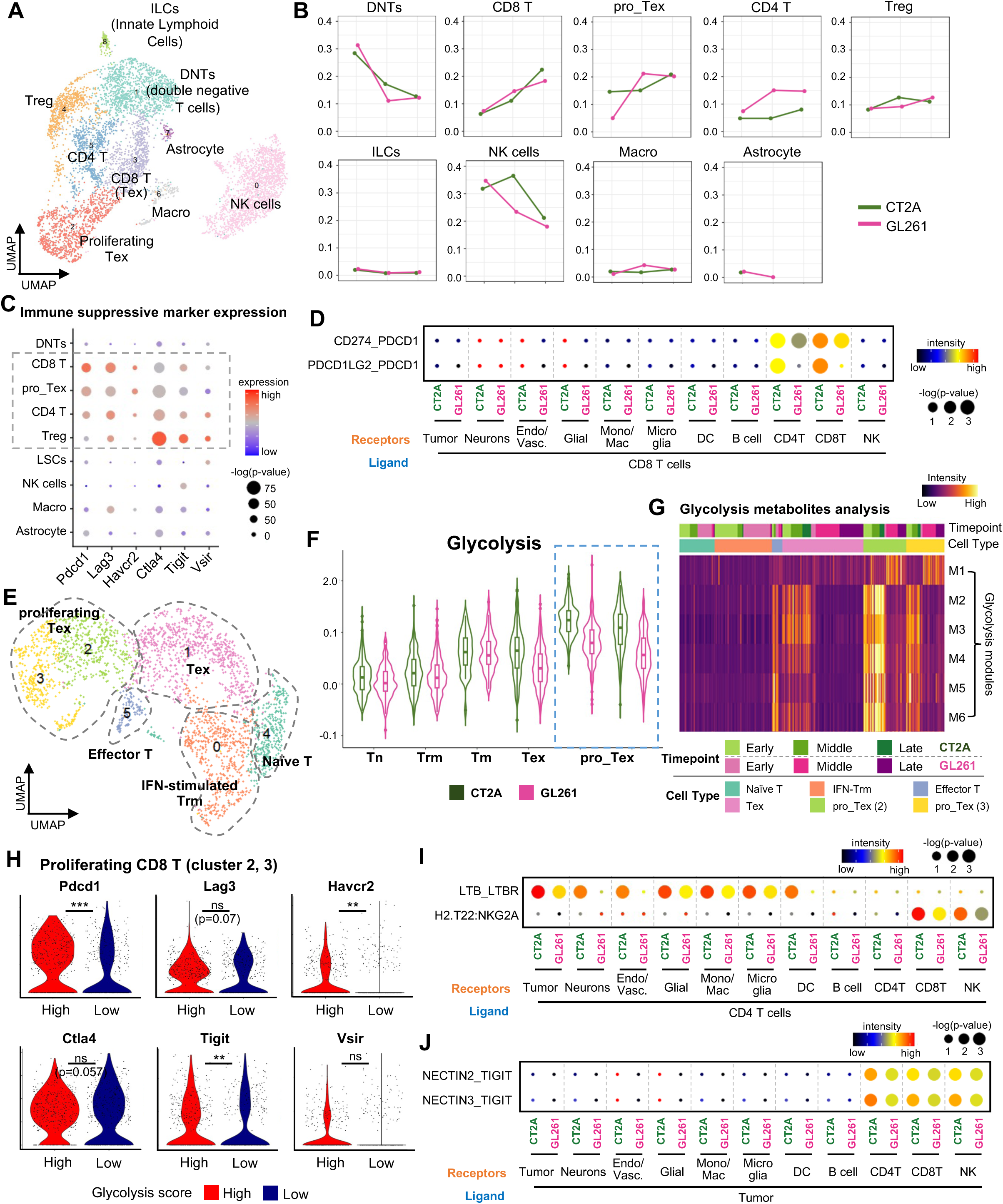
Dynamic changes in lymphocytes and their cell-cell interactions. **A.** UMAP visualization of re-clustered lymphocytes. **B.** Line plot showing changes in lymphocyte population across time points for each cluster. **C.** Dot plot indicating expression levels of representative immune cell dysfunction markers. **D.** Cell-cell interactions in which CD8 T cells act as ligands between the two tumor models in scRNA-seq datasets. Dot size and color represent -log(p-value) and interaction intensity, respectively. **E.** UMAP visualization of re-clustered CD8 T cells. **F.** Violin plot showing glycolysis scores between CT2A and GL261 across CD8 T cell subtypes. **G.** Heatmap indicating module activity of glycolysis metabolites in CD8 T cell subtypes across time points. **H.** Violin plot showing expression levels of dysfunction markers between glycolysis high and low groups in proliferating Tex. **I.** Cell-cell interactions in which CD4 T cells act as ligands between the two tumor models in scRNA-seq datasets. Dot size and color represent -log(p-value) and interaction intensity, respectively. **J.** Cell–cell interactions between tumor and T/NK cells, with cancer cells act as ligands. Dot size and color represent -log(p-value) and interaction intensity, respectively.

We next explored the functional state of CD8+ T cells by sub-clustering them into five functional states (Fig. 6E). While the broad subtype composition was similar between models (Fig. S12A, B), we identified a critical metabolic differentiator: glycolytic activity was elevated in proliferating exhausted T cells (Pro_Tex), particularly in late timepoints from CT2A (Fig. 6F and S12C).

Metabolite flux analysis using scFEA (Alghamdi et al., 2021) further confirmed the accumulation of glycolysis-related metabolites in Pro_Tex from CT2A (Fig. 6G). Interestingly, we observed the upregulation of exhaustion markers, including *Pdcd1*, *Havcr2*, *Tigit*, and *Vsir*, in parallel with enhanced glycolysis (Fig. 6H).

Next, we examined the "receiver" side of the MHC-I signaling identified in earlier sections. In CT2A, CD4+ T cells, CD8+ T cells, and NK cells all exhibited inhibitory interactions (Wang et al., 2022) through the H2-T22:Nkg2a axis (Fig. 6I, Fig. S9D). Inhibitory interactions were observed also via robust *Ltb*:*Ltbr* interactions, which promote tumor progression (Fig. S9D) (Y. Wu et al., 2024). The resulting *Nectin2:Tigit* and *Nectin3:Tigit* interactions likely serve as critical secondary checkpoints (Wienke et al., 2024) that reinforce immune exclusion in late-stage CT2A (Fig. 6J and S9D). Beyond their inhibition via *H2-T22:Nkg2a*, NK cells engaged in *Lgals9*:*P4hb* and *Lgals9*:*Havcr2* signaling, which induce cancer cell migration and T cell exhaustion (Ni et al., 2022; Zhu et al., 2025). Additionally, NK cells in CT2A appear to actively recruit dysfunctional immune cells through *ICAM*:integrin interactions (Fig. S12E,F). Altogether, the CT2A lymphoid landscape is defined by a dense network of upregulated inhibitory interactions beyond simple exhaustion. By co-opting non-classical MHC-I signaling, metabolic stress, and chronic checkpoint activity, the CT2A tumor creates deep immune dysfunction that likely precludes a successful response to immunotherapy.

## Discussion

In this study, we performed multi-omic time courses - including scRNA-seq and ST - on two complementary and widely used murine glioblastoma models, CT2A and GL261. These tumor models respond differently to immune checkpoint blockade and exhibit markedly distinct mortality profiles, suggesting fundamental differences in the tumor and immune compartments that contribute to disease progression. Our data set enabled us to comprehensively investigate the mechanisms driving these divergent outcomes by analyzing the tumor transcriptome, cellular composition of the tumor microenvironment, and interactions among tumor and immune cells.

By mapping our data onto established human GBM meta-modules (Neftel et al., 2019), we show that CT2A and GL261 recapitulate divergent transcriptional states observed in patient tumors. CT2A aligns closely with the Mesenchymal (MES1/2) state, a phenotype historically associated with high-grade malignancy, poor prognosis, and a robust inflammatory niche. In contrast, GL261 resembles the Oligodendrocyte Progenitor (OPC) and Astrocytic (AC) states, which are defined by neuro-developmental programs and a more permissive immune environment. This distinction suggests that these models are equally valid yet complementary tools, each capturing different facets of human GBM biology and treatment resistance. Our analysis indicates that CT2A is transcriptionally heterogeneous, with distinct functional programs identified across multiple cell clusters and support of plasticity in trajectory analysis. In contrast, GL261 concurrently expresses markers associated with MES1/2, OPC, and AC states. GL261 also exhibits a temporal shift toward the OPC state at later stages. These findings suggest that while CT2A represents a mesenchymal model intrinsically heterogeneous and plastic that is, nonetheless, stable over time within the mesenchymal-hypoxia axis, GL261 could be seen as recapitulating the multi-state cellular architecture observed in patient tumors.

Our results also demonstrate that the aggressive CT2A phenotype is driven by a progressive upregulation of hypoxia response and epithelial mesenchymal transition (EMT). These processes are well-established drivers of glioblastoma invasion and therapeutic resistance, resulting in poor prognosis (Iwadate, 2016; Monteiro et al., 2017; Park and Lee, 2022). We then identified three key factors, the thrombospondins (*Thbs1* and *Thbs2*) and osteopontin (*Spp1*), which were co-expressed and correlated with hypoxia response and the EMT pathway within CT2A tumor cell populations. These genes were expressed in a CT2A-specific and time-dependent manner, and mediated numerous computationally inferred tumor-tumor and tumor-immune interactions in CT2A. THBS1 is a secreted matricellular glycoprotein identified as an angiogenic inhibitor (Bornstein, 2009) but also implicated in several pro-tumorigenic processes including cell proliferation and as a pro-invasive gene, namely in GBM (Daubon et al., 2019; Whitehead et al., 2023). Notably, THBS1 has important roles both in the tumor microenvironment (Kaur et al., 2021) and immunity (Kaur and Roberts, 2024). Previous studies have reported that *Thbs1* is either induced under hypoxia (Phelan et al., 1998) or contributes to the establishment of a hypoxic environment, for example, by inhibiting angiogenesis (Bornstein, 2009). The relationship between *Thbs2* and hypoxia is less established, although its function has been studied, showing that *Thbs2* enhances EMT pathways via extracellular matrix remodeling (Zhang et al., 2025) and possibly modulating tumor vasculogenic mimicry (Huang et al., 2023). In our dataset, *Thbs1* was elevated early in CT2A, whereas *Thbs2* expression progressively increased at later CT2A time points. This dynamic expression pattern suggests a potential cooperative mechanism in which *Thbs1* may accelerate hypoxia early in tumor formation, which in turn promotes upregulation of *Thbs2* and subsequent EMT activation at late CT2A. Such a sequential interplay between *Thbs1* and *Thbs2* could reinforce tumor aggressiveness and immune evasion. Importantly, *Thbs1* and *Thbs2* have been associated with poor prognosis across multiple cancer types, and our correlative exploration with survival in human transcriptomic and proteomic datasets confirmed that increased expression of *Thbs1* and *Thbs2* correlates with poor prognosis in GBM, underscoring the translational relevance of our findings. These findings are further supported by spatial transcriptomic mapping of the human GBM ecosystem, which identifies SPP1 as one of the top marker genes defining the Mesenchymal spatial tumor cell state and THBS1/2 as central components of a signaling network facilitating ubiquitous intercellular communication across diverse tissue domains (Khan et al., 2025).

We also identified striking upregulation of MHC Class II (MHC-II) genes in GL261 tumor cells. Our prior work in acute myeloid leukemia (Christopher et al., 2018) and several studies in lung, and colorectal cancers have shown that increased MHC-II expression by tumor cells is associated with improved prognosis and enhanced therapeutic responses by promoting CD8 T cell infiltration and activation (Griffith et al., 2022; Johnson et al., 2020) likely through CD4 T cell help. This is consistent with our prior observations in the GL261 model, where CD4 T cells are required for the efficacy of immune checkpoint blockade (ICI) (Khan et al., 2022).

While some studies associate MHC-II expression in glioblastoma with an aggressive ’myeloid-mimetic’ state (Gangoso et al., 2021) and an immunosuppressive microenvironment, we observed that MHC-II was absent in the CD45-negative fraction of 11 human GBM resections analyzed by flow cytometry. This lack of expression in human samples aligns with the phenotype observed in CT2A, where tumor cells were also refractory to MHC-II induction by IFNγ *in vitro*. Furthermore, we found that *Hmox1*, a representative hypoxia-induced gene, was highly expressed in CT2A. HO-1, encoded by *Hmox1*, was reported to suppress MHC-II expression, providing a mechanistic link between the hypoxic conditions and MHC-II expression (Alaluf et al., 2020; Chora et al., 2007). Consistent with this, pseudotime trajectory analysis revealed that the expression of both classical and non-classical MHC-I genes declined as tumor cells transitioned along Lineage 2 into the hypoxic cluster. Taken together, these findings suggest that hypoxia-driven *Hmox1* may attenuate MHC-II expression in CT2A.

The upregulation of MHC-I genes in CT2A, was intriguing, because the MHC-I machinery is impaired in this model due to deleterious mutations in the peptide-loading machinery, specifically in *Tap1* and *Psmb8* (Iorgulescu et al., 2023). Deficiencies in the Transporter associated with Antigen Processing (TAP) are associated with the presentation of alternative antigens, termed T-cell epitopes associated with impaired peptide processing (TEIPPs), which are loaded onto MHC-I molecules in the absence of canonical peptide transport (Marijt et al., 2018; Van Hall et al., 2006). Given the expression of MHC-I molecules in CT2A and the high exhaustion scores observed in CT2A-infiltrating CD8 T cells we cannot exclude the possibility of antigen recognition, which could occur via TEIPP presentation. Because TEIPP-derived autoantigens are only presented under conditions of impaired processing, they are not subject to central tolerance, unlike most non-mutated self-antigens (Marijt et al., 2018). While we cannot confirm any of these speculations and indeed our data is also consistent with the acquisition of an exhausted state independent of antigen presentation, these observations are nonetheless useful to highlight the possibility that, for personalized glioblastoma treatment, clinical strategies targeting tumors with antigen-presentation deficiencies could consider the exploit of TEIPP-derived epitopes as non-tolerized alternatives to traditional neoantigens.

In addition to cancer cells, we observed striking remodeling of the myeloid populations in both CT2A and GL261. Notably, microglia, the resident brain macrophages with homeostatic functions, progressively lost their canonical identity in CT2A and acquired transcriptional features resembling macrophages. Such a loss of microglial characteristics has also been demonstrated in metastasis-associated microglia, and it has been reported that glioblastoma can influence microglial gene expression to support tumor growth (W. Ma et al., 2024; Maas et al., 2020). Consistent with these observations, we detected pronounced morphological alterations in microglia, especially in CT2A, suggesting that macrophage-like microglial cells facilitate tumor progression and contribute to the establishment of an immunosuppressive niche in CT2A.

Macrophages exhibited robust infiltration and immunosuppression-related interactions in CT2A. Among these, we found that *Spp1*-mediated interactions were correlated with hypoxia and EMT. Previous studies have demonstrated that *Spp1* not only promotes macrophage infiltration but also drives M2 polarization, contributing to poor prognosis and therapeutic resistance across multiple cancer types (Song et al., 2023; Tan et al., 2025; Wang et al., 2024). These findings suggest that hypoxia within CT2A activates *Spp1*-mediated interactions, which, in turn, promote macrophage polarization and TAM infiltration, ultimately reinforcing immune suppression and poor therapeutic responsiveness. In addition, we observed enriched interactions with *Ccl7*, *Ccl2*, and *Cxcl14* that promote recruitment of TAM and reinforce immune suppression at late CT2A (Goto et al., 2023; Gschwandtner et al., 2019; Khan et al., 2024; X. Wu et al., 2024). The recruited macrophages promote glioblastoma proliferation, and survival by producing LDHA-containing extracellular vesicles (Khan et al., 2024).

Functional analysis of these macrophages revealed divergent metabolic and hypoxic trajectories. In CT2A, hypoxia scores in myeloid cells increased throughout tumor progression, whereas these scores decreased in the GL261 model. This sustained hypoxic environment in CT2A correlated with elevated glycolytic markers in both myeloid cells and proliferating exhausted T cells. These data indicate that the CT2A microenvironment imposes a metabolic state characterized by high glycolytic flux, which is associated with the terminal exhaustion of infiltrating CD8 T cells. In addition to metabolic stress, myeloid-driven inhibitory checkpoint interactions, including *Lgals9:Havcr2* and *Cd274:Pdcd1*, exacerbated T cell exhaustion and reinforced an immunosuppressive niche (He and Xu, 2020; Jin et al., 2023; Kandel et al., 2021). Collectively, these findings suggest that while GL261 maintains an inflammatory state permissive for T-cell activity, the CT2A microenvironment establishes a sustained hypoxic and metabolic barrier that facilitates tumor progression and immune dysfunction.

We also identified in CT2A Pdcd1- and Tigit-dependent interactions, potentially contributing to T-cell exhaustion reinforced by NK cell–mediated inhibitory signaling, collectively sustaining dysfunctional immune infiltration despite stable lymphocyte abundance.

In this study, we propose that in CT2A a complex network composed of hypoxia, *Thbs*/*Spp1*-mediated interactions, microglial plasticity, macrophage infiltration, and lymphocyte dysfunction orchestrate tumor progression. This network establishes immunosuppressive microenvironment that not only promotes tumor progression but also underlies the immune suppression. By contrast, GL261 tumors, while aggressive, retain features of inflammatory signaling and less extensive immunosuppressive characteristics, which may account for their improved prognosis.

Our study has several limitations. First, while CT2A and GL261 are the most widely used murine GBM models and provide valuable insight into human disease, neither is a perfect model for human preclinical GBM. Second, our observations are based primarily on transcriptomic and spatial analyses, and functional validation will be necessary to identify direct causal relationships between observed pathways and tumor progression. Nevertheless, the association between high thrombospondin and osteopontin expression and poor survival was also detected in public human datasets, and key findings such as lack of MHC-II expression were further validated by flow cytometry, supporting our results.

Our comprehensive analysis of time-resolved single-cell and spatial transcriptomic data demonstrated that hypoxia-related transcriptional changes, thrombospondin- and osteopontin-mediated interactions, decreased MHC Class II gene expression, and complex alterations across immune cells may contribute to the pro-tumorigenic, immunosuppressive microenvironment of CT2A. These insights highlight critical biological differences between CT2A and GL261, and between different human tumor cell states, and lay the foundation for potential therapeutic strategies to improve immunotherapy efficacy in glioblastoma.

## Methods

### Tumor growth time courses

Mice were injected with 5 x10^6^ tumor cells (*n*=5 for GL261 and CT2A) into the right frontal lobe of female C57/Bl6 mice. Axial and coronal T1 (post-gadolinium contrast) and T2 MRI were obtained using a small animal adaptor on post-injection days 5, 7, 10, 12, 14, 17, and 19, or until mice met pre-determined survival endpoints. Segmentation and volumetric analyses were performed in ITK-SNAP.

### Sample collection

At all three time points, mice (*n=5*) were perfused, tumors were harvested, processed into a single cell suspension (myelin removal, ACK lysis, dead cell removal) and processed for single cell RNA sequencing. At the latter time points (days 14 and 17), tumors were harvested, fixed in formalin, and paraffin-embedded for spatial transcriptomics (in process, 10x Visium FFPE kit).

### Single-cell RNA-seq library construction and analysis

We used the 10x Genomics 5’ V2 Chromium Gene Expression Solution to generate cDNA libraries for each time point. Libraries were sequenced on an Illumina Novaseq S2 flow cell to a target depth of 50,000 reads/spot. Sequencing reads were demultiplexed, aligned, and counted using the Cellranger 5.0 pipeline. Downstream processing steps, including filtering, doublet removal, normalization, scaling, identification of variable genes, dimensionality reduction, and clustering were performed in Seurat.

### Visium data library construction and analysis

Each tumor model was sampled for Visium analysis at the middle and late time points determined by the MRI growth curves. Each section (i.e. sample) was preserved in FFPE, and cDNA libraries were generated according to the Visium Spatial Gene Expression for FFPE User Guide Rev. E. All libraries were sequenced on an Illumina Novaseq S2 flow cell to a target depth of 50,000 reads/spot and yielded high quality transcriptome data. Demultiplexing, image and transcript alignment, and transcript quantification was performed using the Spaceranger pipeline (10x Genomics, v1.2, mm10 reference). Initial downstream analysis - including normalization for per-spot sequencing depth using the SCTransform algorithm, principal component analysis, UMAP layout, graph-based clustering, differential gene expression analysis, and spatially variable gene identification - was performed using the R package Seurat (V4). Previously defined gene expression modules associated with tumor and immune cell states were incorporated into this analysis using the AddModuleScore Seurat function. Spatially variable biological processes and cell types were also inferred using clusterProfileR.

### Cell-cell interaction analysis

CellChat was used to perform the cell-cell communication analysis (Jin et al., 2025). To prioritize the most supported cell-cell interactions, we integrated analyses from CellChat and CellPhoneDB databases (Troulé et al., 2025). Since CellPhoneDB is designed for human gene references, mouse gene lists were converted to their human orthologs using g:Profiler (Reimand et al., 2007). Normalized UMI count and cluster information were used as the input file for statistical analysis function with p-value 0.05. Visualization was done using ggplot2 R package (v3.5.2). In addition, Nichenet was used to identify cell-to-cell interactions and targets that were regulated by ligands (Browaeys et al., 2020). Their interaction and target regulating potential were calculated utilizing the Nichent database, and it was visualized by the pheatmap R package (v1.0.12).

### Gene co-expression network

Gene co-expression networks were constructed using the high-dimensional Weighted Gene Co-expression Network Analysis (hdWGCNA) R package (v0.4.06) (Morabito et al., 2023). The parameter settings were configured in accordance with the manufacturer’s instructions. The calculation of the weighted adjacency between genes was based on the Pearson correlation. To enhance the accuracy of the co-expression network, a soft power threshold of 6 was set. This threshold facilitated the removal of noise and weak connections, contributing to the construction of an accurate co-expression network. Each network was characterized by its biological pathways using Enrichr (https://maayanlab.cloud/Enrichr/) (Kuleshov et al., 2016). The differences in fold change were visualized using the pheatmap R package.

### Biological functions and correlation

The hypoxia, epithelial-mesenchymal transition (EMT), and glycolysis score was calculated using the AddModuleScore function in Seurat (Hao et al., 2021). The genes comprising each module were from the MSigDB hallmark database (Subramanian et al., 2005). The biological pathways in the heatmap were identified using Enrichr, which is a gene ontology webserver interface. Correlation between biological pathway score (hypoxia and EMT) with genes was calculated by Pearson correlation. The correlation was calculated using our normalized expression matrix and the cor function on the R package stats (v4.5.0) was used for analysis.

### Kaplan meier curve

Kaplan meier curve was screenshot from PROTEIN ATLAS web-interface (https://www.proteinatlas.org/) (Uhlén et al., 2015). We applied the best expression cut off option and p-value provide by PROTEIN ATLAS.

### Metabolic flux analysis

The scFEA packages was employed to estimate metabolites flux in CD8 T cells from scRNA-seq (Alghamdi et al., 2021). Normalized expression matrix was used as input, with the modules database for each metabolite provided by a scFEA. scFEA predicts metabolism flux based on each metabolite variation using neural network model. The results were visualized as heatmap, where positive intensity indicates metabolite accumulation and negative values reveal outflow of metabolite.

## References

Alaluf, E., Vokaer, B., Detavernier, A., Azouz, A., Splittgerber, M., Carrette, A., Boon, L., Libert, F., Soares, M.P., Le Moine, A., Goriely, S., 2020. Heme oxygenase-1 orchestrates the immunosuppressive program of tumor-associated macrophages. JCI Insight. 10.1172/jci.insight.133929

Alghamdi, N., Chang, W., Dang, P., Lu, X., Wan, C., Gampala, S., Huang, Z., Wang, J., Ma, Q., Zang, Y., Fishel, M., Cao, S., Zhang, C., 2021. A graph neural network model to estimate cell-wise metabolic flux using single-cell RNA-seq data. Genome Res. 31, 1867–1884. 10.1101/gr.271205.120

Alspach, E., Lussier, D.M., Miceli, A.P., Kizhvatov, I., DuPage, M., Luoma, A.M., Meng, W., Lichti, C.F., Esaulova, E., Vomund, A.N., Runci, D., Ward, J.P., Gubin, M.M., Medrano, R.F.V., Arthur, C.D., White, J.M., Sheehan, K.C.F., Chen, A., Wucherpfennig, K.W., Jacks, T., Unanue, E.R., Artyomov, M.N., Schreiber, R.D., 2019. MHC-II neoantigens shape tumour immunity and response to immunotherapy. Nature 574, 696–701. 10.1038/s41586-019-1671-8

Ausman, J.I., Shapiro, W.R., Rall, D.P., 1970. Studies on the chemotherapy of experimental brain tumors: development of an experimental model. Cancer Res 30, 2394–2400.

Bied, M., Ho, W.W., Ginhoux, F., Blériot, C., 2023. Roles of macrophages in tumor development: a spatiotemporal perspective. Cell Mol Immunol 20, 983–992. 10.1038/s41423-023-01061-6

Bornstein, P., 2009. Thrombospondins function as regulators of angiogenesis. J. Cell Commun. Signal. 3, 189–200. 10.1007/s12079-009-0060-8

Browaeys, R., Saelens, W., Saeys, Y., 2020. NicheNet: modeling intercellular communication by linking ligands to target genes. Nat Methods 17, 159–162. 10.1038/s41592-019-0667-5

Candolfi, M., Curtin, J.F., Nichols, W.S., Muhammad, A.G., King, G.D., Pluhar, G.E., McNiel, E.A., Ohlfest, J.R., Freese, A.B., Moore, P.F., Lerner, J., Lowenstein, P.R., Castro, M.G., 2007. Intracranial glioblastoma models in preclinical neuro-oncology: neuropathological characterization and tumor progression. J Neurooncol 85, 133–148. 10.1007/s11060-007-9400-9

Chauvin, J.-M., Zarour, H.M., 2020. TIGIT in cancer immunotherapy. J Immunother Cancer 8, e000957. 10.1136/jitc-2020-000957

Chen, J., 2024. TNFSF12 is associated with breast cancer prognosis and immune cell infiltration. Am J Transl Res 16, 4120–4133. 10.62347/IDTK3218

Chen, Z., Lau, K.S., 2023. Advances in Mapping Tumor Progression from Precancer Atlases. Cancer Prevention Research 16, 439–447. 10.1158/1940-6207.CAPR-22-0473

Chora, Â.A., Fontoura, P., Cunha, A., Pais, T.F., Cardoso, S., Ho, P.P., Lee, L.Y., Sobel, R.A., Steinman, L., Soares, M.P., 2007. Heme oxygenase–1 and carbon monoxide suppress autoimmune neuroinflammation. J. Clin. Invest. 117, 438–447. 10.1172/JCI28844

Christofides, A., Strauss, L., Yeo, A., Cao, C., Charest, A., Boussiotis, V.A., 2022. The complex role of tumor-infiltrating macrophages. Nat Immunol 23, 1148–1156. 10.1038/s41590-022-01267-2

Christopher, M.J., Petti, A.A., Rettig, M.P., Miller, C.A., Chendamarai, E., Duncavage, E.J., Klco, J.M., Helton, N.M., O’Laughlin, M., Fronick, C.C., Fulton, R.S., Wilson, R.K., Wartman, L.D., Welch, J.S., Heath, S.E., Baty, J.D., Payton, J.E., Graubert, T.A., Link, D.C., Walter, M.J., Westervelt, P., Ley, T.J., DiPersio, J.F., 2018. Immune Escape of Relapsed AML Cells after Allogeneic Transplantation. New England Journal of Medicine 379, 2330–2341. 10.1056/NEJMoa1808777

Comba, A., Faisal, S.M., Dunn, P.J., Argento, A.E., Hollon, T.C., Al-Holou, W.N., Varela, M.L., Zamler, D.B., Quass, G.L., Apostolides, P.F., Abel, C., Brown, C.E., Kish, P.E., Kahana, A., Kleer, C.G., Motsch, S., Castro, M.G., Lowenstein, P.R., 2022. Spatiotemporal analysis of glioma heterogeneity reveals COL1A1 as an actionable target to disrupt tumor progression. Nat Commun 13, 3606. 10.1038/s41467-022-31340-1

Culp, P.A., Choi, D., Zhang, Y., Yin, J., Seto, P., Ybarra, S.E., Su, M., Sho, M., Steinle, R., Wong, M.H.L., Evangelista, F., Grove, J., Cardenas, M., James, M., Hsi, E.D., Chao, D.T., Powers, D.B., Ramakrishnan, V., Dubridge, R., 2010. Antibodies to TWEAK Receptor Inhibit Human Tumor Growth through Dual Mechanisms. Clinical Cancer Research 16, 497–508. 10.1158/1078-0432.CCR-09-1929

Daubon, T., Léon, C., Clarke, K., Andrique, L., Salabert, L., Darbo, E., Pineau, R., Guérit, S., Maitre, M., Dedieu, S., Jeanne, A., Bailly, S., Feige, J.-J., Miletic, H., Rossi, M., Bello, L., Falciani, F., Bjerkvig, R., Bikfalvi, A., 2019. Deciphering the complex role of thrombospondin-1 in glioblastoma development. Nat Commun 10, 1146. 10.1038/s41467-019-08480-y

Fan, Y., Potdar, A.A., Gong, Y., Eswarappa, S.M., Donnola, S., Lathia, J.D., Hambardzumyan, D., Rich, J.N., Fox, P.L., 2014. Profilin-1 phosphorylation directs angiocrine expression and glioblastoma progression through HIF-1α accumulation. Nat Cell Biol 16, 445–456. 10.1038/ncb2954

Fleetwood, A.J., Achuthan, A., Schultz, H., Nansen, A., Almholt, K., Usher, P., Hamilton, J.A., 2014. Urokinase Plasminogen Activator Is a Central Regulator of Macrophage Three-Dimensional Invasion, Matrix Degradation, and Adhesion. The Journal of Immunology 192, 3540–3547. 10.4049/jimmunol.1302864

Gangoso, E., Southgate, B., Bradley, L., Rus, S., Galvez-Cancino, F., McGivern, N., Güç, E., Kapourani, C.-A., Byron, A., Ferguson, K.M., Alfazema, N., Morrison, G., Grant, V., Blin, C., Sou, I., Marques-Torrejon, M.A., Conde, L., Parrinello, S., Herrero, J., Beck, S., Brandner, S., Brennan, P.M., Bertone, P., Pollard, J.W., Quezada, S.A., Sproul, D., Frame, M.C., Serrels, A., Pollard, S.M., 2021. Glioblastomas acquire myeloid-affiliated transcriptional programs via epigenetic immunoediting to elicit immune evasion. Cell 184, 2454–2470.e26. 10.1016/j.cell.2021.03.023

Gatta, A., Shaik, A.K.B., Shallak, M., Chiaravalli, A.M., Cerati, M., La Rosa, S., Accolla, R.S., Forlani, G., 2025. CIITA-modified glioblastomas vaccinate and induce cross-protection against heterologous wild-type glioblastomas. J Transl Med 23. 10.1186/s12967-025-06816-5

Gau, D., Daoud, A., Allen, A., Joy, M., Sagan, A., Lee, S., Lucas, P.C., Duensing, S., Boone, D., Osmanbeyoglu, H.U., Roy, P., 2023. Vascular endothelial profilin-1 drives a protumorigenic tumor microenvironment and tumor progression in renal cancer. Journal of Biological Chemistry 299, 105044. 10.1016/j.jbc.2023.105044

Genoud, V., Marinari, E., Nikolaev, S.I., Castle, J.C., Bukur, V., Dietrich, P.-Y., Okada, H., Walker, P.R., 2018. Responsiveness to anti-PD-1 and anti-CTLA-4 immune checkpoint blockade in SB28 and GL261 mouse glioma models. Oncoimmunology 7, e1501137. 10.1080/2162402X.2018.1501137

Ghoochani, A., Schwarz, M.A., Yakubov, E., Engelhorn, T., Doerfler, A., Buchfelder, M., Bucala, R., Savaskan, N.E., Eyüpoglu, I.Y., 2016. MIF-CD74 signaling impedes microglial M1 polarization and facilitates brain tumorigenesis. Oncogene 35, 6246–6261. 10.1038/onc.2016.160

Goto, A., Komura, S., Kato, K., Maki, R., Hirakawa, A., Tomita, H., Hirata, A., Yamada, Y., Akiyama, H., 2023. C-X-C domain ligand 14-mediated stromal cell–macrophage interaction as a therapeutic target for hand dermal fibrosis. Commun Biol 6, 1173. 10.1038/s42003-023-05558-8

Griffith, B.D., Turcotte, S., Lazarus, J., Lima, F., Bell, S., Delrosario, L., McGue, J., Krishnan, S., Oneka, M.D., Nathan, H., Smith, J.J., D’Angelica, M.I., Shia, J., Di Magliano, M.P., Rao, A., Frankel, T.L., 2022. MHC Class II Expression Influences the Composition and Distribution of Immune Cells in the Metastatic Colorectal Cancer Microenvironment. Cancers 14, 4092. 10.3390/cancers14174092

Gschwandtner, M., Derler, R., Midwood, K.S., 2019. More Than Just Attractive: How CCL2 Influences Myeloid Cell Behavior Beyond Chemotaxis. Front. Immunol. 10, 2759. 10.3389/fimmu.2019.02759

Guo, K.S., Brodsky, A.S., 2023. Tumor collagens predict genetic features and patient outcomes. npj Genom. Med. 8, 15. 10.1038/s41525-023-00358-9

Haddad, A.F., Young, J.S., Amara, D., Berger, M.S., Raleigh, D.R., Aghi, M.K., Butowski, N.A., 2021. Mouse models of glioblastoma for the evaluation of novel therapeutic strategies. Neurooncol Adv 3, vdab100. 10.1093/noajnl/vdab100

Hao, Y., Hao, S., Andersen-Nissen, E., Mauck, W.M., Zheng, S., Butler, A., Lee, M.J., Wilk, A.J., Darby, C., Zager, M., Hoffman, P., Stoeckius, M., Papalexi, E., Mimitou, E.P., Jain, J., Srivastava, A., Stuart, T., Fleming, L.M., Yeung, B., Rogers, A.J., McElrath, J.M., Blish, C.A., Gottardo, R., Smibert, P., Satija, R., 2021. Integrated analysis of multimodal single-cell data. Cell 184, 3573–3587.e29. 10.1016/j.cell.2021.04.048

He, X., Xu, C., 2020. Immune checkpoint signaling and cancer immunotherapy. Cell Res 30, 660– 669. 10.1038/s41422-020-0343-4

Huang, J., Wang, C., Hou, Y., Tian, Y., Li, Y., Zhang, H., Zhang, L., Li, W., 2023. Molecular mechanisms of Thrombospondin-2 modulates tumor vasculogenic mimicry by PI3K/AKT/mTOR signaling pathway. Biomedicine & Pharmacotherapy 167, 115455. 10.1016/j.biopha.2023.115455

Iorgulescu, J.B., Ruthen, N., Ahn, R., Panagioti, E., Gokhale, P.C., Neagu, M., Speranza, M.C., Eschle, B.K., Soroko, K.M., Piranlioglu, R., Datta, M., Krishnan, S., Yates, K.B., Baker, G.J., Jain, R.K., Suvà, M.L., Neuberg, D., White, F.M., Chiocca, E.A., Freeman, G.J., Sharpe, A.H., Wu, C.J., Reardon, D.A., 2023. Antigen presentation deficiency, mesenchymal differentiation, and resistance to immunotherapy in the murine syngeneic CT2A tumor model. Front. Immunol. 14, 1297932. 10.3389/fimmu.2023.1297932

Iwadate, Y., 2016. Epithelial-mesenchymal transition in glioblastoma progression. Oncology Letters 11, 1615–1620. 10.3892/ol.2016.4113

Jin, S., Plikus, M.V., Nie, Q., 2025. CellChat for systematic analysis of cell–cell communication from single-cell transcriptomics. Nat Protoc 20, 180–219. 10.1038/s41596-024-01045-4

Jin, X., Cai, Y., Xue, G., Que, J., Cheng, R., Yang, Y., Xiao, L., Lin, X., Xu, C., Wang, P., Xu, Z., Nie, H., Jiang, Q., 2023. Identification of shared characteristics in tumor-infiltrating T cells across 15 cancers. Molecular Therapy - Nucleic Acids 32, 189–202. 10.1016/j.omtn.2023.03.007

Johnson, A.M., Bullock, B.L., Neuwelt, A.J., Poczobutt, J.M., Kaspar, R.E., Li, H.Y., Kwak, J.W., Hopp, K., Weiser-Evans, M.C.M., Heasley, L.E., Schenk, E.L., Clambey, E.T., Nemenoff, R.A., 2020. Cancer Cell–Intrinsic Expression of MHC Class II Regulates the Immune Microenvironment and Response to Anti–PD-1 Therapy in Lung Adenocarcinoma. The Journal of Immunology 204, 2295–2307. 10.4049/jimmunol.1900778

Kandel, S., Adhikary, P., Li, G., Cheng, K., 2021. The TIM3/Gal9 signaling pathway: An emerging target for cancer immunotherapy. Cancer Letters 510, 67–78. 10.1016/j.canlet.2021.04.011

Kang, M., Gulati, G.S., Brown, E.L., Qi, Z., Avagyan, S., Armenteros, J.J.A., Gleyzer, R., Zhang, W., Steen, C.B., D’Silva, J.P., Schwab, J., Clarke, M.F., Chaudhuri, A.A., Newman, A.M., 2025. Improved reconstruction of single-cell developmental potential with CytoTRACE 2. Nat Methods 22, 2258–2263. 10.1038/s41592-025-02857-2

Katzenelenbogen, Y., Sheban, F., Yalin, A., Yofe, I., Svetlichnyy, D., Jaitin, D.A., Bornstein, C., Moshe, A., Keren-Shaul, H., Cohen, M., Wang, S.-Y., Li, B., David, E., Salame, T.-M., Weiner, A., Amit, I., 2020. Coupled scRNA-Seq and Intracellular Protein Activity Reveal an Immunosuppressive Role of TREM2 in Cancer. Cell 182, 872–885.e19. 10.1016/j.cell.2020.06.032

Kaur, S., Bronson, S.M., Pal-Nath, D., Miller, T.W., Soto-Pantoja, D.R., Roberts, D.D., 2021. Functions of Thrombospondin-1 in the Tumor Microenvironment. Int J Mol Sci 22, 4570. 10.3390/ijms22094570

Kaur, S., Roberts, D.D., 2024. Emerging functions of thrombospondin-1 in immunity. Seminars in Cell & Developmental Biology, Thrombospondin proteins 155, 22–31. 10.1016/j.semcdb.2023.05.008

Khan, F., Lin, Y., Ali, H., Pang, L., Dunterman, M., Hsu, W.-H., Frenis, K., Grant Rowe, R., Wainwright, D.A., McCortney, K., Billingham, L.K., Miska, J., Horbinski, C., Lesniak, M.S., Chen, P., 2024. Lactate dehydrogenase A regulates tumor-macrophage symbiosis to promote glioblastoma progression. Nat Commun 15, 1987. 10.1038/s41467-024-46193-z

Khan, S.M., Desai, R., Coxon, A., Livingstone, A., Dunn, G.P., Petti, A., Johanns, T.M., 2022. Impact of CD4 T cells on intratumoral CD8 T-cell exhaustion and responsiveness to PD-1 blockade therapy in mouse brain tumors. J Immunother Cancer 10, e005293. 10.1136/jitc-2022-005293

Khan, S.M., Wang, A.Z., Desai, R.R., McCornack, C.R., Sun, R., Dahiya, S.M., Foltz, J.A., Sherpa, N.D., Leavitt, L., West, T., Wang, A.F., Krbanjevic, A., Choi, B.D., Leuthardt, E.C., Patel, B., Charest, A., Kim, A.H., Dunn, G.P., Petti, A.A., 2025. Mapping the spatial architecture of glioblastoma from core to edge delineates niche-specific tumor cell states and intercellular interactions. 10.1101/2025.04.04.647096

Kirishima, M., Yokoyama, S., Akahane, T., Higa, N., Uchida, H., Yonezawa, H., Matsuo, K., Yamamoto, J., Yoshimoto, K., Hanaya, R., Tanimoto, A., 2024. Prognosis prediction via histological evaluation of cellular heterogeneity in glioblastoma. Sci Rep 14, 24955. 10.1038/s41598-024-76826-8

Kochan, G., Escors, D., Breckpot, K., Guerrero-Setas, D., 2013. Role of non-classical MHC class I molecules in cancer immunosuppression. OncoImmunology 2, e26491. 10.4161/onci.26491

Kuleshov, M.V., Jones, M.R., Rouillard, A.D., Fernandez, N.F., Duan, Q., Wang, Z., Koplev, S., Jenkins, S.L., Jagodnik, K.M., Lachmann, A., McDermott, M.G., Monteiro, C.D., Gundersen, G.W., Ma’ayan, A., 2016. Enrichr: a comprehensive gene set enrichment analysis web server 2016 update. Nucleic Acids Res 44, W90–W97. 10.1093/nar/gkw377

La Manno, G., Siletti, K., Furlan, A., Gyllborg, D., Vinsland, E., Mossi Albiach, A., Mattsson Langseth, C., Khven, I., Lederer, A.R., Dratva, L.M., Johnsson, A., Nilsson, M., Lönnerberg, P., Linnarsson, S., 2021. Molecular architecture of the developing mouse brain. Nature 596, 92–96. 10.1038/s41586-021-03775-x

Lee, P.J., Jiang, B.-H., Chin, B.Y., Iyer, N.V., Alam, J., Semenza, G.L., Choi, AugustineM.K., 1997. Hypoxia-inducible Factor-1 Mediates Transcriptional Activation of the Heme Oxygenase-1 Gene in Response to Hypoxia. Journal of Biological Chemistry 272, 5375–5381. 10.1074/jbc.272.9.5375

Li, R., Hebert, J.D., Lee, T.A., Xing, H., Boussommier-Calleja, A., Hynes, R.O., Lauffenburger, D.A., Kamm, R.D., 2017. Macrophage-Secreted TNFα and TGFβ1 Influence Migration Speed and Persistence of Cancer Cells in 3D Tissue Culture via Independent Pathways. Cancer Research 77, 279–290. 10.1158/0008-5472.CAN-16-0442

Liu, Connor J, Schaettler, M., Blaha, D.T., Bowman-Kirigin, J.A., Kobayashi, D.K., Livingstone, A.J., Bender, D., Miller, C.A., Kranz, D.M., Johanns, T.M., Dunn, G.P., 2020a. Treatment of an aggressive orthotopic murine glioblastoma model with combination checkpoint blockade and a multivalent neoantigen vaccine. Neuro Oncol 22, 1276–1288. 10.1093/neuonc/noaa050

Liu, Connor J., Schaettler, M., Blaha, D.T., Bowman-Kirigin, J.A., Kobayashi, D.K., Livingstone, A.J., Bender, D., Miller, C.A., Kranz, D.M., Johanns, T.M., Dunn, G.P., 2020. Treatment of an aggressive orthotopic murine glioblastoma model with combination checkpoint blockade and a multivalent neoantigen vaccine. Neuro Oncol 22, 1276–1288. 10.1093/neuonc/noaa050

Liu, Connor J, Schaettler, M., Blaha, D.T., Bowman-Kirigin, J.A., Kobayashi, D.K., Livingstone, A.J., Bender, D., Miller, C.A., Kranz, D.M., Johanns, T.M., Dunn, G.P., 2020b. Treatment of an aggressive orthotopic murine glioblastoma model with combination checkpoint blockade and a multivalent neoantigen vaccine. Neuro-Oncology 22, 1276–1288. 10.1093/neuonc/noaa050

Ma, C., Chen, J., Ji, J., Zheng, Y., Liu, Y., Wang, J., Chen, T., Chen, H., Chen, Z., Zhou, Q., Hou, C., Ke, Y., 2024. Therapeutic modulation of APP-CD74 axis can activate phagocytosis of TAMs in GBM. Biochimica et Biophysica Acta (BBA) - Molecular Basis of Disease 1870, 167449. 10.1016/j.bbadis.2024.167449

Ma, W., Oswald, J., Rios Angulo, A., Chen, Q., 2024. Tmem119 expression is downregulated in a subset of brain metastasis-associated microglia. BMC Neurosci 25, 6. 10.1186/s12868-024-00846-3

Maas, S.L.N., Abels, E.R., Van De Haar, L.L., Zhang, X., Morsett, L., Sil, S., Guedes, J., Sen, P., Prabhakar, S., Hickman, S.E., Lai, C.P., Ting, D.T., Breakefield, X.O., Broekman, M.L.D., El Khoury, J., 2020. Glioblastoma hijacks microglial gene expression to support tumor growth. J Neuroinflammation 17, 120. 10.1186/s12974-020-01797-2

Marijt, K.A., Blijleven, L., Verdegaal, E.M.E., Kester, M.G., Kowalewski, D.J., Rammensee, H.-G., Stevanović, S., Heemskerk, M.H.M., Van Der Burg, S.H., Van Hall, T., 2018. Identification of non-mutated neoantigens presented by TAP-deficient tumors. Journal of Experimental Medicine 215, 2325–2337. 10.1084/jem.20180577

Marin, I.A., Gutman-Wei, A.Y., Chew, K.S., Raissi, A.J., Djurisic, M., Shatz, C.J., 2022. The nonclassical MHC class I Qa-1 expressed in layer 6 neurons regulates activity-dependent plasticity via microglial CD94/NKG2 in the cortex. Proc. Natl. Acad. Sci. U.S.A. 119, e2203965119. 10.1073/pnas.2203965119

Martínez-Murillo, R., 2007. Standardization of an orthotopic mouse brain tumor model following transplantation of CT-2A astrocytoma cells. Histology and Histopathology 1309–1326. 10.14670/HH-22.1309

Mikolajewicz, N., Tatari, N., Wei, J., Savage, N., Granda Farias, A., Dimitrov, V., Chen, D., Zador, Z., Dasgupta, K., Aguilera-Uribe, M., Xiao, Y.-X., Lee, S.Y., Mero, P., McKenna, D., Venugopal, C., Brown, K.R., Han, H., Singh, S., Moffat, J., 2024. Functional profiling of murine glioma models highlights targetable immune evasion phenotypes. Acta Neuropathol 148, 74. 10.1007/s00401-024-02831-w

Miller, T.E., El Farran, C.A., Couturier, C.P., Chen, Z., D’Antonio, J.P., Verga, J., Villanueva, M.A., Gonzalez Castro, L.N., Tong, Y.E., Saadi, T.A., Chiocca, A.N., Zhang, Y., Fischer, D.S., Heiland, D.H., Guerriero, J.L., Petrecca, K., Suva, M.L., Shalek, A.K., Bernstein, B.E., 2025. Programs, origins and immunomodulatory functions of myeloid cells in glioma. Nature 1–11. 10.1038/s41586-025-08633-8

Monteiro, A., Hill, R., Pilkington, G., Madureira, P., 2017. The Role of Hypoxia in Glioblastoma Invasion. Cells 6, 45. 10.3390/cells6040045

Morabito, S., Reese, F., Rahimzadeh, N., Miyoshi, E., Swarup, V., 2023. hdWGCNA identifies co-expression networks in high-dimensional transcriptomics data. Cell Reports Methods 3, 100498. 10.1016/j.crmeth.2023.100498

Neftel, C., Laffy, J., Filbin, M.G., Hara, T., Shore, M.E., Rahme, G.J., Richman, A.R., Silverbush, D., Shaw, M.L., Hebert, C.M., Dewitt, J., Gritsch, S., Perez, E.M., Gonzalez Castro, L.N., Lan, X., Druck, N., Rodman, C., Dionne, D., Kaplan, A., Bertalan, M.S., Small, J., Pelton, K., Becker, S., Bonal, D., Nguyen, Q.-D., Servis, R.L., Fung, J.M., Mylvaganam, R., Mayr, L., Gojo, J., Haberler, C., Geyeregger, R., Czech, T., Slavc, I., Nahed, B.V., Curry, W.T., Carter, B.S., Wakimoto, H., Brastianos, P.K., Batchelor, T.T., Stemmer-Rachamimov, A., Martinez-Lage, M., Frosch, M.P., Stamenkovic, I., Riggi, N., Rheinbay, E., Monje, M., Rozenblatt-Rosen, O., Cahill, D.P., Patel, A.P., Hunter, T., Verma, I.M., Ligon, K.L., Louis, D.N., Regev, A., Bernstein, B.E., Tirosh, I., Suvà, M.L., 2019. An Integrative Model of Cellular States, Plasticity, and Genetics for Glioblastoma. Cell 178, 835–849.e21. 10.1016/j.cell.2019.06.024

Newcomb, E.W., Zagzag, D., 2009. The Murine GL261 Glioma Experimental Model to Assess Novel Brain Tumor Treatments, in: Meir, E.G. (Ed.), CNS Cancer: Models, Markers, Prognostic Factors, Targets, and Therapeutic Approaches. Humana Press, Totowa, NJ, pp. 227–241. 10.1007/978-1-60327-553-8_12

Ni, X., Wu, W., Sun, X., Ma, J., Yu, Z., He, X., Cheng, J., Xu, P., Liu, H., Shang, T., Xi, S., Wang, J., Zhang, J., Chen, Z., 2022. Interrogating glioma-M2 macrophage interactions identifies Gal-9/Tim-3 as a viable target against PTEN-null glioblastoma. Sci. Adv. 8, eabl5165. 10.1126/sciadv.abl5165

Oh, T., Fakurnejad, S., Sayegh, E.T., Clark, A.J., Ivan, M.E., Sun, M.Z., Safaee, M., Bloch, O., James, C.D., Parsa, A.T., 2014. Immunocompetent murine models for the study of glioblastoma immunotherapy. J Transl Med 12, 107. 10.1186/1479-5876-12-107

Park, J.H., Lee, H.K., 2022. Current Understanding of Hypoxia in Glioblastoma Multiforme and Its Response to Immunotherapy. Cancers 14, 1176. 10.3390/cancers14051176

Parker, N.R., Khong, P., Parkinson, J.F., Howell, V.M., Wheeler, H.R., 2015. Molecular Heterogeneity in Glioblastoma: Potential Clinical Implications. Front. Oncol. 5. 10.3389/fonc.2015.00055

Patel, A.P., Tirosh, I., Trombetta, J.J., Shalek, A.K., Gillespie, S.M., Wakimoto, H., Cahill, D.P., Nahed, B.V., Curry, W.T., Martuza, R.L., Louis, D.N., Rozenblatt-Rosen, O., Suvà, M.L., Regev, A., Bernstein, B.E., 2014. Single-cell RNA-seq highlights intratumoral heterogeneity in primary glioblastoma. Science 344, 1396–1401. 10.1126/science.1254257

Phelan, M.W., Forman, L.W., Perrine, S.P., Faller, D.V., 1998. Hypoxia increases thrombospondin-1 transcript and protein in cultured endothelial cells. Journal of Laboratory and Clinical Medicine 132, 519–529. 10.1016/S0022-2143(98)90131-7

Qiang, L., Wu, T., Zhang, H.-W., Lu, N., Hu, R., Wang, Y.-J., Zhao, L., Chen, F.-H., Wang, X.-T., You, Q.-D., Guo, Q.-L., 2012. HIF-1α is critical for hypoxia-mediated maintenance of glioblastoma stem cells by activating Notch signaling pathway. Cell Death Differ 19, 284–294. 10.1038/cdd.2011.95

Raredon, M.S.B., Yang, J., Kothapalli, N., Lewis, W., Kaminski, N., Niklason, L.E., Kluger, Y., 2023. Comprehensive visualization of cell–cell interactions in single-cell and spatial transcriptomics with NICHES. Bioinformatics 39, btac775. 10.1093/bioinformatics/btac775

Reardon, D.A., Gokhale, P.C., Klein, S.R., Ligon, K.L., Rodig, S.J., Ramkissoon, S.H., Jones, K.L., Conway, A.S., Liao, X., Zhou, J., Wen, P.Y., Van Den Abbeele, A.D., Hodi, F.S., Qin, L., Kohl, N.E., Sharpe, A.H., Dranoff, G., Freeman, G.J., 2016. Glioblastoma Eradication Following Immune Checkpoint Blockade in an Orthotopic, Immunocompetent Model. Cancer Immunology Research 4, 124–135. 10.1158/2326-6066.CIR-15-0151

Reimand, J., Kull, M., Peterson, H., Hansen, J., Vilo, J., 2007. g:Profiler—a web-based toolset for functional profiling of gene lists from large-scale experiments. Nucleic Acids Research 35, W193–W200. 10.1093/nar/gkm226

Saelens, W., Cannoodt, R., Todorov, H., Saeys, Y., 2019. A comparison of single-cell trajectory inference methods. Nat Biotechnol 37, 547–554. 10.1038/s41587-019-0071-9

Schoppmeyer, R., Zhao, R., Cheng, H., Hamed, M., Liu, C., Zhou, X., Schwarz, E.C., Zhou, Y., Knörck, A., Schwär, G., Ji, S., Liu, L., Long, J., Helms, V., Hoth, M., Yu, X., Qu, B., 2017. Human profilin 1 is a negative regulator of CTL mediated cell-killing and migration. Eur J Immunol 47, 1562–1572. 10.1002/eji.201747124

Senner, V., Ratzinger, S., Mertsch, S., Grässel, S., Paulus, W., 2008. Collagen XVI expression is upregulated in glioblastomas and promotes tumor cell adhesion. FEBS Letters 582, 3293–3300. 10.1016/j.febslet.2008.09.017

Song, J., Sun, J., Jing, S., Zhang, T., Wang, J., Liu, Y., 2023. A high level of secreted phosphoprotein 1 is associated with macrophage infiltration and poor prognosis in hepatocellular carcinoma. iLIVER 2, 26–35. 10.1016/j.iliver.2023.01.004

Street, K., Risso, D., Fletcher, R.B., Das, D., Ngai, J., Yosef, N., Purdom, E., Dudoit, S., 2018. Slingshot: cell lineage and pseudotime inference for single-cell transcriptomics. BMC Genomics 19, 477. 10.1186/s12864-018-4772-0

Subramanian, A., Tamayo, P., Mootha, V.K., Mukherjee, S., Ebert, B.L., Gillette, M.A., Paulovich, A., Pomeroy, S.L., Golub, T.R., Lander, E.S., Mesirov, J.P., 2005. Gene set enrichment analysis: A knowledge-based approach for interpreting genome-wide expression profiles. Proc. Natl. Acad. Sci. U.S.A. 102, 15545–15550. 10.1073/pnas.0506580102

Szatmári, T., Lumniczky, K., Désaknai, S., Trajcevski, S., Hídvégi, E.J., Hamada, H., Sáfrány, G., 2006. Detailed characterization of the mouse glioma 261 tumor model for experimental glioblastoma therapy. Cancer Sci 97, 546–553. 10.1111/j.1349-7006.2006.00208.x

Tan, Y., Yang, Y.-G., Zhang, X., Zhao, L., Wang, X., Liu, W., 2025. Tumor cell-derived osteopontin promotes tumor fibrosis indirectly via tumor-associated macrophages. J Transl Med 23, 432. 10.1186/s12967-025-06444-z

Troulé, K., Petryszak, R., Cakir, B., Cranley, J., Harasty, A., Prete, M., Tuong, Z.K., Teichmann, S.A., Garcia-Alonso, L., Vento-Tormo, R., 2025. CellPhoneDB v5: inferring cell–cell communication from single-cell multiomics data. Nat Protoc 1–29. 10.1038/s41596-024-01137-1

Uhlén, M., Fagerberg, L., Hallström, B.M., Lindskog, C., Oksvold, P., Mardinoglu, A., Sivertsson, Å., Kampf, C., Sjöstedt, E., Asplund, A., Olsson, I., Edlund, K., Lundberg, E., Navani, S., Szigyarto, C.A.-K., Odeberg, J., Djureinovic, D., Takanen, J.O., Hober, S., Alm, T., Edqvist, P.-H., Berling, H., Tegel, H., Mulder, J., Rockberg, J., Nilsson, P., Schwenk, J.M., Hamsten, M., Von Feilitzen, K., Forsberg, M., Persson, L., Johansson, F., Zwahlen, M., Von Heijne, G., Nielsen, J., Pontén, F., 2015. Tissue-based map of the human proteome. Science 347, 1260419. 10.1126/science.1260419

Uhlen, M., Zhang, C., Lee, S., Sjöstedt, E., Fagerberg, L., Bidkhori, G., Benfeitas, R., Arif, M., Liu, Z., Edfors, F., Sanli, K., Von Feilitzen, K., Oksvold, P., Lundberg, E., Hober, S., Nilsson, P., Mattsson, J., Schwenk, J.M., Brunnström, H., Glimelius, B., Sjöblom, T., Edqvist, P.-H., Djureinovic, D., Micke, P., Lindskog, C., Mardinoglu, A., Ponten, F., 2017. A pathology atlas of the human cancer transcriptome. Science 357, eaan2507. 10.1126/science.aan2507

Urosevic, M., Dummer, R., 2008. Human Leukocyte Antigen–G and Cancer Immunoediting. Cancer Research 68, 627–630. 10.1158/0008-5472.CAN-07-2704

Van Hall, T., Wolpert, E.Z., Van Veelen, P., Laban, S., Van Der Veer, M., Roseboom, M., Bres, S., Grufman, P., De Ru, A., Meiring, H., De Jong, A., Franken, K., Teixeira, A., Valentijn, R., Drijfhout, J.W., Koning, F., Camps, M., Ossendorp, F., Kärre, K., Ljunggren, H.-G., Melief, C.J.M., Offringa, R., 2006. Selective cytotoxic T-lymphocyte targeting of tumor immune escape variants. Nat Med 12, 417–424. 10.1038/nm1381

Wang, C., Li, Y., Wang, L., Han, Y., Gao, X., Li, T., Liu, M., Dai, L., Du, R., 2024. SPP1 represents a therapeutic target that promotes the progression of oesophageal squamous cell carcinoma by driving M2 macrophage infiltration. Br J Cancer 130, 1770–1782. 10.1038/s41416-024-02683-x

Wang, X., Xiong, H., Ning, Z., 2022. Implications of NKG2A in immunity and immune-mediated diseases. Front. Immunol. 13, 960852. 10.3389/fimmu.2022.960852

Wei, Q., Singh, O., Ekinci, C., Gill, J., Li, M., Mamatjan, Y., Karimi, S., Bunda, S., Mansouri, S., Aldape, K., Zadeh, G., 2021. TNFα secreted by glioma associated macrophages promotes endothelial activation and resistance against anti-angiogenic therapy. acta neuropathol commun 9, 67. 10.1186/s40478-021-01163-0

Wei, R., He, S., Bai, S., Sei, E., Hu, M., Thompson, A., Chen, K., Krishnamurthy, S., Navin, N.E., 2022. Spatial charting of single-cell transcriptomes in tissues. Nat Biotechnol 40, 1190– 1199. 10.1038/s41587-022-01233-1

Whitehead, C.A., Morokoff, A.P., Kaye, A.H., Drummond, K.J., Mantamadiotis, T., Stylli, S.S., 2023. Invadopodia associated Thrombospondin-1 contributes to a post-therapy pro-invasive response in glioblastoma cells. Experimental Cell Research 431, 113743. 10.1016/j.yexcr.2023.113743

Wienke, J., Visser, L.L., Kholosy, W.M., Keller, K.M., Barisa, M., Poon, E., Munnings-Tomes, S., Himsworth, C., Calton, E., Rodriguez, A., Bernardi, R., Van Den Ham, F., Van Hooff, S.R., Matser, Y.A.H., Tas, M.L., Langenberg, K.P.S., Lijnzaad, P., Borst, A.L., Zappa, E., Bergsma, F.J., Strijker, J.G.M., Verhoeven, B.M., Mei, S., Kramdi, A., Restuadi, R., Sanchez-Bernabeu, A., Cornel, A.M., Holstege, F.C.P., Gray, J.C., Tytgat, G.A.M., Scheijde-Vermeulen, M.A., Wijnen, M.H.W.A., Dierselhuis, M.P., Straathof, K., Behjati, S., Wu, W., Heck, A.J.R., Koster, J., Nierkens, S., Janoueix-Lerosey, I., De Krijger, R.R., Baryawno, N., Chesler, L., Anderson, J., Caron, H.N., Margaritis, T., Van Noesel, M.M., Molenaar, J.J., 2024. Integrative analysis of neuroblastoma by single-cell RNA sequencing identifies the NECTIN2-TIGIT axis as a target for immunotherapy. Cancer Cell 42, 283–300.e8. 10.1016/j.ccell.2023.12.008

Wu, A., Oh, S., Wiesner, S.M., Ericson, K., Chen, L., Hall, W.A., Champoux, P.E., Low, W.C., Ohlfest, J.R., 2008. Persistence of CD133^+^ Cells in Human and Mouse Glioma Cell Lines: Detailed Characterization of GL261 Glioma Cells with Cancer Stem Cell-Like Properties. Stem Cells and Development 17, 173–184. 10.1089/scd.2007.0133

Wu, X., Mi, T., Jin, L., Ren, C., Wang, J., Zhang, Z., Liu, J., Wang, Z., Guo, P., He, D., 2024. Dual roles of HK3 in regulating the network between tumor cells and tumor-associated macrophages in neuroblastoma. Cancer Immunol Immunother 73, 122. 10.1007/s00262-024-03702-9

Wu, Y., Zhao, S., Guo, W., Liu, Y., Requena Mullor, M.D.M., Rodrìguez, R.A., Wei, R., 2024. Systematic analysis of the prognostic value and immunological function of LTBR in human cancer. Aging. 10.18632/aging.205356

Yeo, A.T., Rawal, S., Delcuze, B., Christofides, A., Atayde, A., Strauss, L., Balaj, L., Rogers, V.A., Uhlmann, E.J., Varma, H., Carter, B.S., Boussiotis, V.A., Charest, A., 2022. Single-cell RNA sequencing reveals evolution of immune landscape during glioblastoma progression. Nat Immunol 23, 971–984. 10.1038/s41590-022-01215-0

Yin, W., Zhu, H., Tan, J., Xin, Z., Zhou, Q., Cao, Y., Wu, Z., Wang, L., Zhao, M., Jiang, X., Ren, C., Tang, G., 2021. Identification of collagen genes related to immune infiltration and epithelial-mesenchymal transition in glioma. Cancer Cell Int 21, 276. 10.1186/s12935-021-01982-0

Yun, S.P., Ryu, J.M., Jang, M.W., Han, H.J., 2011. Interaction of profilin-1 and F-actin via a β-arrestin-1/JNK signaling pathway involved in prostaglandin E2 -induced human mesenchymal stem cells migration and proliferation. Journal Cellular Physiology 226, 559–571. 10.1002/jcp.22366

Zarodniuk, M., Steele, A., Lu, X., Li, J., Datta, M., 2023. CNS tumor stroma transcriptomics identify perivascular fibroblasts as predictors of immunotherapy resistance in glioblastoma patients. npj Genom. Med. 8, 35. 10.1038/s41525-023-00381-w

Zhang, L.-P., Zhang, Z.-G., Guan, J., Li, L.-Q., 2025. Thrombospondin 2 drives liver metastasis in skin cutaneous melanoma via regulation of angiogenesis and extracellular matrix remodeling. Melanoma Research 35, 306–316. 10.1097/CMR.0000000000001055

Zhu, X., Zhang, Y., Yu, A., Xiao, X., 2025. Myeloid-Derived LGALS9 - P4HB Immune Interaction Promotes Metastasis in Gastric Cancer Through Enhanced Cell Proliferation and Lipid Metabolism. J Cellular Molecular Medi 29, e70661. 10.1111/jcmm.70661

Zimmerman, H.M., Arnold, H., 1941. Experimental Brain Tumors. I. Tumors Produced with Methylcholanthrene*. Cancer Research 1, 919–938.

