## Supplemental Information for "Integrated spatial and single-cell transcriptomic analysis of aggressive glioblastoma growth dynamics"

**Supplementary Note 1:** Longitudinal Tumor Growth Dynamics and Sampling

Aiming at identifying comparable time points for sampling, we utilized serial Magnetic Resonance Imaging (MRI) to generate growth curves for each model (Fig. 1B). The results showed that, while the two models grew at similar rates as detected by MRI, CT2A-bearing mice reached clinical endpoint sooner. Timepoints were thus selected to represent developmental stage and terminal point, rather than chronological age or tumor volume. This resulted in a delayed collection schedule for GL261. Consistent with the rationale leading to this decision, we consistently recovered fewer cells from the GL261 model for the same selected stage despite delayed collection. At the early timepoint, we observed significantly fewer tumor cells in GL261 (*N=19* cells) compared to CT2A (*N=2184* - Fig. 2C, Fig. S1A,B), even after pooling 5 animals per model. By the late stage in ST samples, the ratio of tumor cells over neurons was 85% in CT2A compared to 34% in GL261, even though CT2A being collected two days earlier (Fig. S1C).

**Supplementary Note 2:** Technical and Algorithm Justifications

To ensure robust analysis of the longitudinal single-cell and spatial data, several state-of-the-art computational tools were utilized.

Spatial Deconvolution: Visium ST data was deconvolved using CellTrek (1), a machine learning approach that maps cells from a scRNA-seq reference to their most likely location in the tissue slice. To provide a baseline for healthy brain architecture, we spiked our CT2A and GL261 scRNA-seq data with normal adolescent mouse brain snRNA-seq data (2).

Trajectory Analysis: Pseudo-temporal trajectories in CT2A tumor cells were modeled using the Slingshot (3) algorithm, which is recognized as one of the top-ranked methods for identifying branched developmental architectures in single-cell data (4).

Potency Inference: Cellular differentiation states were assessed via CytoTRACE2 (5), an approach which applies a deep learning framework to infer cellular potency based on transcriptional diversity.

Gene Co-expression: To identify integrated biological programs, we used hdWGCNA (high-dimensional Weighted Gene Co-expression Network Analysis) (6) to identify modules of co-expressed genes, specifically identifying mTOR, HIF1A, and Wnt/beta-catenin modules in CT2A.

**Supplementary Note 3:** Shared Intercellular Communication and ECM Architecture.

Beyond the model-specific interactions highlighted in the main text, CellChat (7) identified a conserved set of shared ligand-receptor (L-R) interactions. Heatmaps showing extended lists of interactions initiated by the tumor (tumor as “sender”) are presented in Fig. S5, as a resource. For maximized functional interpretation, we present these results for a high-resolution annotation of our single cell dataset. In the following text, we highlight some of the take-way messages that can be drawn, with focus on what could be common features of these two models.

*Conserved Myeloid and Neuronal Signaling, and Vascular Co-option*

Both models prioritized tumor-to-myeloid signaling through the App/Mif–Cd74 and Trem2–Tyrobp axes, pathways previously shown to induce immunosuppressive myeloid phenotypes (Fig. S5A,D – annotations in black) (8–10). These conserved immunosuppressive pathways are potentially bolstered in both models by the expression of additional inhibitory and immune checkpoint-like ligands, such as Galectin-9 (Lgals9)(11–13), with tumor potentially communicating via Lgals9 with both myeloid and lymphoid cells (Fig. S5A-C). Interestingly, we also detected a relatively strong tumor-tumor interaction between Lgals9 and P4HB in both models, which could contribute to enhanced tumor proliferation and EMT (14) (Fig. S5D). Additionally, both models utilized neuronal programs such as synaptic adhesion molecules (Ncam1, Tenm3/4) and guidance cues (Sema6a, Ptprs). This shared core extends to the perivascular niche, potentially coordinating vascular co-option and myeloid recruitment via Fn1, Hspg2, and Jam3 (Fig. S5 E,F).

*Conserved ECM Remodeling but Divergent Utilization of Collagen Types*

ECM remodeling was highly enriched in overrepresentation analysis when the tumor cells of both models were compared (Fig 2A,D). While a shared feature of both models, the molecular components were highly model-specific. CT2A upregulated basement membrane and pro-invasive collagens (Col3a1, Col4a1/2, Col5a1/2, Col6a1/2, and Col8a1). These collagen types have been linked to EMT and immunosuppression in GBM (15,16). In contrast, GL261 upregulated chondrogenic-like ECM programs, which are associated with developmental and lineage programs (Col2a1, Col9a1/2/3, and Col11a1/2), along with Col16a1, which has been associated with tumor adhesion in GBM (17). Immunofluorescence (IF) staining confirmed these differences: CT2A tumors were densely packed with COL3A1 depositions organized into linear, spiraling fibers, while in GL261 COL9A2 formed distinct, punctuated, and diffuse depositions (Fig. S6A).
