## Supplemental Figures for "Integrated spatial and single-cell transcriptomic analysis of aggressive glioblastoma growth dynamics"

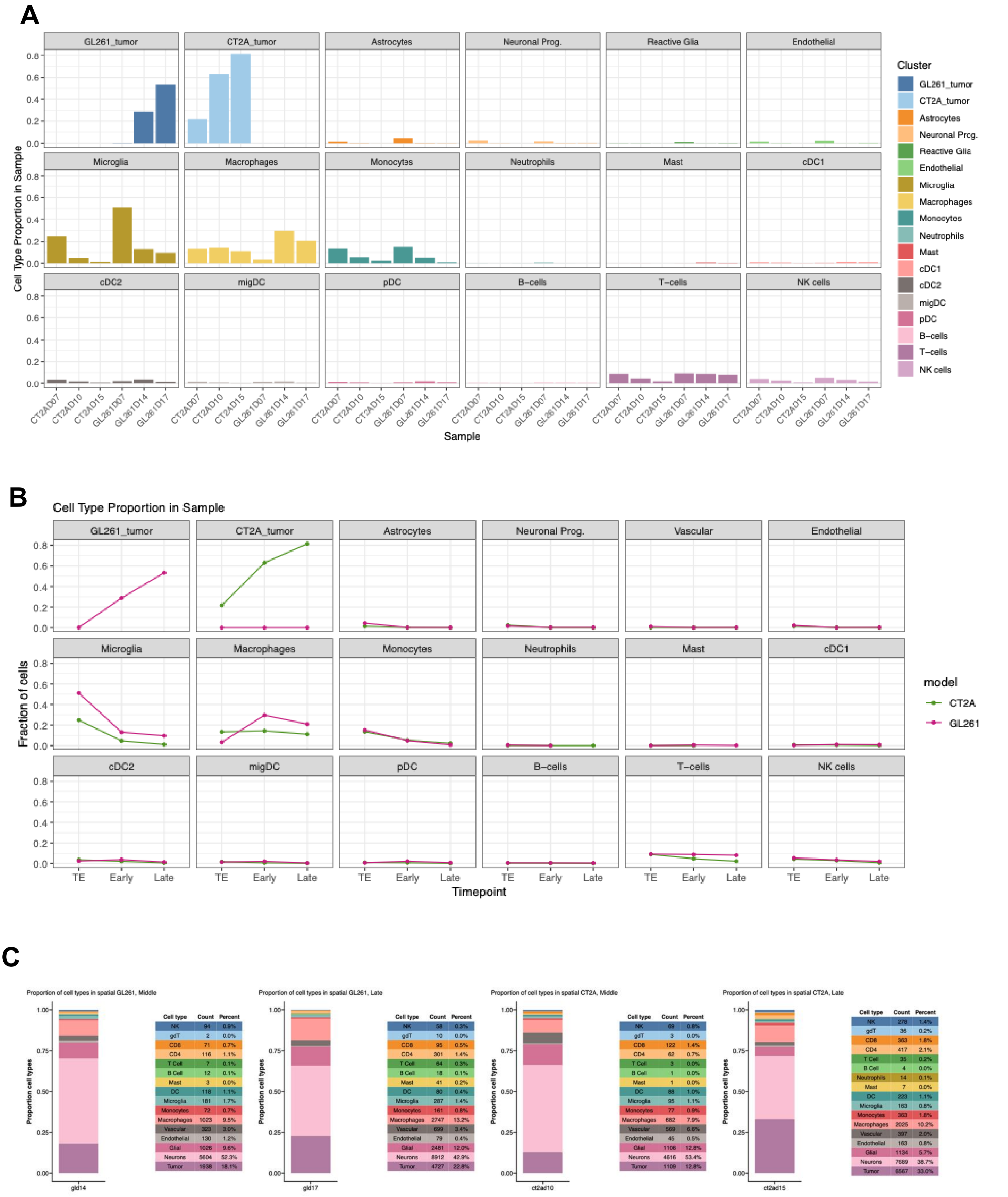


Proportion of cell types in spatial GL261, Middle

| Cell type | Count | Percent |
|---|---|---|
| NK | 94 | 0.9% |
| gdT | 2 | 0.0% |
| CD8 | 71 | 0.7% |
| CD4 | 116 | 1.1% |
| T Cell | 7 | 0.1% |
| B Cell | 12 | 0.1% |
| Mast | 3 | 0.0% |
| DC | 118 | 1.1% |
| Microglia | 181 | 1.7% |
| Monocytes | 72 | 0.7% |
| Macrophages | 1023 | 9.5% |
| Vascular | 323 | 3.0% |
| Endothelial | 130 | 1.2% |
| Glial | 1026 | 9.6% |
| Neurons | 5604 | 52.3% |
| Tumor | 1938 | 18.1% |

Proportion of cell types in spatial GL261, Late

| Cell type | Count | Percent |
|---|---|---|
| NK | 58 | 0.3% |
| gdT | 10 | 0.0% |
| CD8 | 95 | 0.5% |
| CD4 | 301 | 1.4% |
| T Cell | 64 | 0.3% |
| B Cell | 18 | 0.1% |
| Mast | 41 | 0.2% |
| DC | 80 | 0.4% |
| Microglia | 287 | 1.4% |
| Monocytes | 161 | 0.8% |
| Macrophages | 2747 | 13.2% |
| Vascular | 699 | 3.4% |
| Endothelial | 79 | 0.4% |
| Glial | 2481 | 12.0% |
| Neurons | 8912 | 42.9% |
| Tumor | 4727 | 22.8% |

Proportion of cell types in spatial CT2A, Middle

| Cell type | Count | Percent |
|---|---|---|
| NK | 69 | 0.8% |
| gdT | 3 | 0.0% |
| CD8 | 122 | 1.4% |
| CD4 | 62 | 0.7% |
| T Cell | 3 | 0.0% |
| B Cell | 1 | 0.0% |
| Mast | 1 | 0.0% |
| DC | 88 | 1.0% |
| Microglia | 95 | 1.1% |
| Monocytes | 77 | 0.9% |
| Macrophages | 682 | 7.9% |
| Vascular | 569 | 6.6% |
| Endothelial | 45 | 0.5% |
| Glial | 1106 | 12.8% |
| Neurons | 4616 | 53.4% |
| Tumor | 1109 | 12.8% |

Proportion of cell types in spatial CT2A, Late

| Cell type | Count | Percent |
|---|---|---|
| NK | 278 | 1.4% |
| gdT | 36 | 0.2% |
| CD8 | 363 | 1.8% |
| CD4 | 417 | 2.1% |
| T Cell | 35 | 0.2% |
| B Cell | 4 | 0.0% |
| Neutrophils | 14 | 0.1% |
| Mast | 7 | 0.0% |
| DC | 223 | 1.1% |
| Microglia | 163 | 0.8% |
| Monocytes | 363 | 1.8% |
| Macrophages | 2025 | 10.2% |
| Vascular | 397 | 2.0% |
| Endothelial | 163 | 0.8% |
| Glial | 1134 | 5.7% |
| Neurons | 7689 | 38.7% |
| Tumor | 6567 | 33.0% |

**Supplementary figure1. Overall cell-type distribution across timepoints in the two models.**

**A, B.** (A) Bar plot and (B) line plot showing changes in cell type proportion across different samples and time in the entire sample. The cell populations were identified by unbiased clustering and a minimal baseline of GL261 was clustering together with the tumor CT2A cluster. **C.** Proportions and cell counts from projected single cell cells types onto the spatial transcriptomics.

**Supplementary Table 1. Comparative transcriptional programs in CT2A and GL261 tumor cells:** example of top differentially expressed genes (DEGs) highlighting distinct biological pathways in each model (related to Fig. 2A).

| Biological Program | CT2A | | GL261 | |
|---|---|---|---|---|
| | **Signature** | **Representative genes** | **Signature** | **Representative genes** |
| **Core tumor identity** | Mesenchymal / stromal-like | Spp1, Bgn, S100a4, Prrx2, Fosl1, Cav1 | Neuroglial / oligodendrocyte-like | Sox10, Olig1, Olig2, Cnp, Bcas1, Ptprz1 |
| **ECM / matrix organization** | Fibrotic, invasive ECM remodeling | Col3a1, Col4a1, Col5a1, Col6a1, Thbs1, Thbs2, Mmp14, Bgn, Serpinh1 | Developmental / lineage-associated ECM | Col9a1, Col9a2, Col11a1, Col11a2, Col16a1 |
| **Invasion / migration** | EMT-like motility | Mmp14, Spp1, Vcam1, Edil3, Itgb5, Actn1, Myof | Invasion | Mcam, Lgals3, Fat1, Trio, Pcdh17 |
| **Hippo-YAP/TAZ signaling** | Active transcriptional output | Wwtr1, Tead2 | - | - |
| **Developmental / lineage programs** | Mesenchymal patterning / morphogenesis | Hoxc10, Sox9, Nr2f2, Prrx2, Vax2 | Neural / oligodendrocyte lineage | Sox10, Sox6, Olig1, Olig2, Plagl1, Lbh |
| **Angiogenesis / vascular interaction** | Vascular remodeling | Vcam1, Col4a1, Edil3, Cav1, Igfbp3, F2r | - | Mcam |
| **Immune / inflammatory signaling** | Wound-healing / stromal inflammation | Spp1, Serpinf1, Anxa1, Gpx8, Pla2g7 | Antigen presentation + IFN-like | Cd74, H2-Aa, H2-Ab1, H2-Eb1, H2-DMa, C4b, Ifi27l2a |
| **Antigen presentation** | MHC-I non-classical | H2-Q6, H2-Q7 | MHC-II | Cd74, H2-Aa, H2-Ab1, H2-Eb1, H2-DMa |
| **Neuroglial differentiation / myelination** | - | - | - | Sox10, Olig1, Olig2, Cnp, Bcas1, Gjc3, S100b |
| **Stress / metabolic adaptation** | Oxidative stress / injury response | Gstm1, Sgk1, Nupr1, Apod | Metabolic support / survival | Ckb, Moxd1, Asrgl1, Crip1 |
| **Tumor-promoting modulators** | ECM-driven invasion | Thbs1, Thbs2, Mmp14, Spp1 | Immune modulation / survival | Lgals3, Gpnmb, Serpina3n |

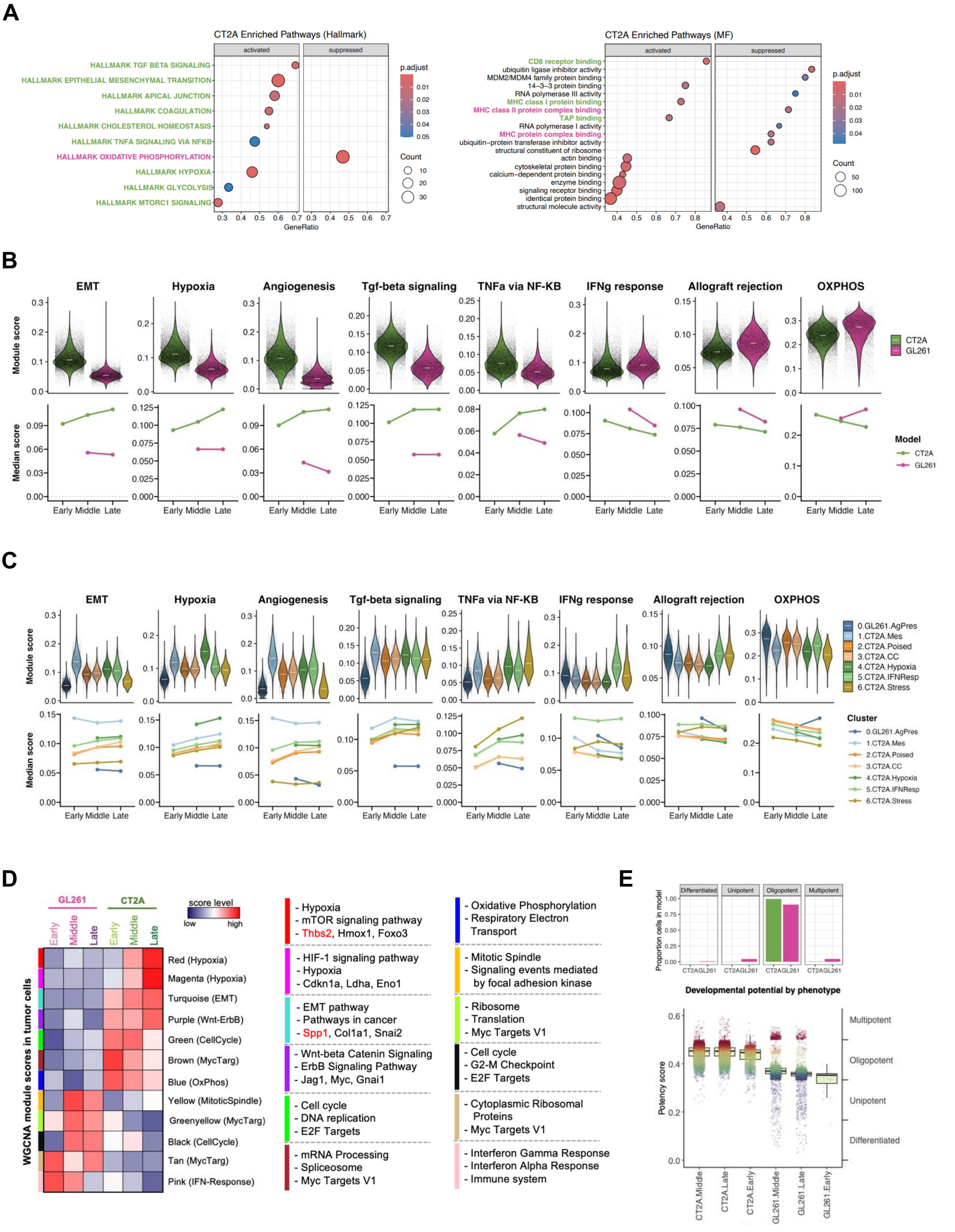
A
CT2A Enriched Pathways (Hallmark)
activated
suppressed
HALLMARK TGF BETA SIGNALING
HALLMARK EPITHELIAL MESENCHYMAL TRANSITION
HALLMARK APICAL JUNCTION
HALLMARK COAGULATION
HALLMARK CHOLESTEROL HOMEOSTASIS
HALLMARK TNFA SIGNALING VIA NFKB
HALLMARK OXIDATIVE PHOSPHORYLATION
HALLMARK HYPOXIA
HALLMARK GLYCOLYSIS
HALLMARK MTORC1 SIGNALING
GeneRatio
p.adjust
Count
CT2A Enriched Pathways (MF)
CD8 receptor binding
ubiquitin ligase inhibitor activity
MDM2/MDM4 family protein binding
14-3-3 protein binding
RNA polymerase III activity
MHC class I protein binding
MHC class II protein complex binding
TAP binding
RNA polymerase I activity
MHC protein complex binding
ubiquitin-protein transferase inhibitor activity
structural constituent of ribosome
actin binding
cytoskeletal protein binding
calcium-dependent protein binding
enzyme binding
signaling receptor binding
identical protein binding
structural molecule activity
B
EMT
Hypoxia
Angiogenesis
Tgf-beta signaling
TNFa via NF-KB
IFNg response
Allograft rejection
OXPHOS
Module score
Median score
Early Middle Late
CT2A
GL261
Model
C
0.GL261.AgPres
1.CT2A.Mes
2.CT2A.Poised
3.CT2A.CC
4.CT2A.Hypoxia
5.CT2A.IFNResp
6.CT2A.Stress
Cluster
D
GL261
CT2A
Early
Middle
Late
score level
low
high
WGCNA module scores in tumor cells
Red (Hypoxia)
Magenta (Hypoxia)
Turquoise (EMT)
Purple (Wnt-ErbB)
Green (CellCycle)
Brown (MycTarg)
Blue (OxPhos)
Yellow (MitoticSpindle)
Greenyellow (MycTarg)
Black (CellCycle)
Tan (MycTarg)
Pink (IFN-Response)
- Hypoxia
- mTOR signaling pathway
- Thbs2, Hmox1, Foxo3
- HIF-1 signaling pathway
- Hypoxia
- Cdkn1a, Ldha, Eno1
- EMT pathway
- Pathways in cancer
- Spp1, Col1a1, Snai2
- Wnt-beta Catenin Signaling
- ErbB Signaling Pathway
- Jag1, Myc, Gnai1
- Cell cycle
- DNA replication
- E2F Targets
- mRNA Processing
- Spliceosome
- Myc Targets V1
- Oxidative Phosphorylation
- Respiratory Electron Transport
- Mitotic Spindle
- Signaling events mediated by focal adhesion kinase
- Ribosome
- Translation
- Myc Targets V1
- Cell cycle
- G2-M Checkpoint
- E2F Targets
- Cytoplasmic Ribosomal Proteins
- Myc Targets V1
- Interferon Gamma Response
- Interferon Alpha Response
- Immune system
E
Differentiated
Unipotent
Oligopotent
Multipotent
Proportion cells in model
CT2AGL261
Developmental potential by phenotype
Potency score
Multipotent
Oligopotent
Unipotent
Differentiated
CT2A.Middle
CT2A.Late
CT2A.Early
GL261.Middle
GL261.Late
GL261.Early

**Supplementary Figure S2. Comparative transcriptomic and stemness profiling of CT2A and GL261 models.**

**(A)** Gene Set Enrichment Analysis (GSEA) of biological pathways based on Hallmark and Molecular Function (MF) gene sets. **(B)** Expression levels of Hallmark signature scores (from MSigDB database) in tumor cells in the two models (top; violin plots) and along longitudinal timepoints (bottom; linepots). **(C)** Same as (B) grouped by tumor cluster. **(D)** Functional annotation and representative hub genes for each hdWGCNA co-expression module. **(E)** Assessment of cellular stemness using CytoTRACE2, presented as categorical distributions (top) and median scores across timepoints (bottom).

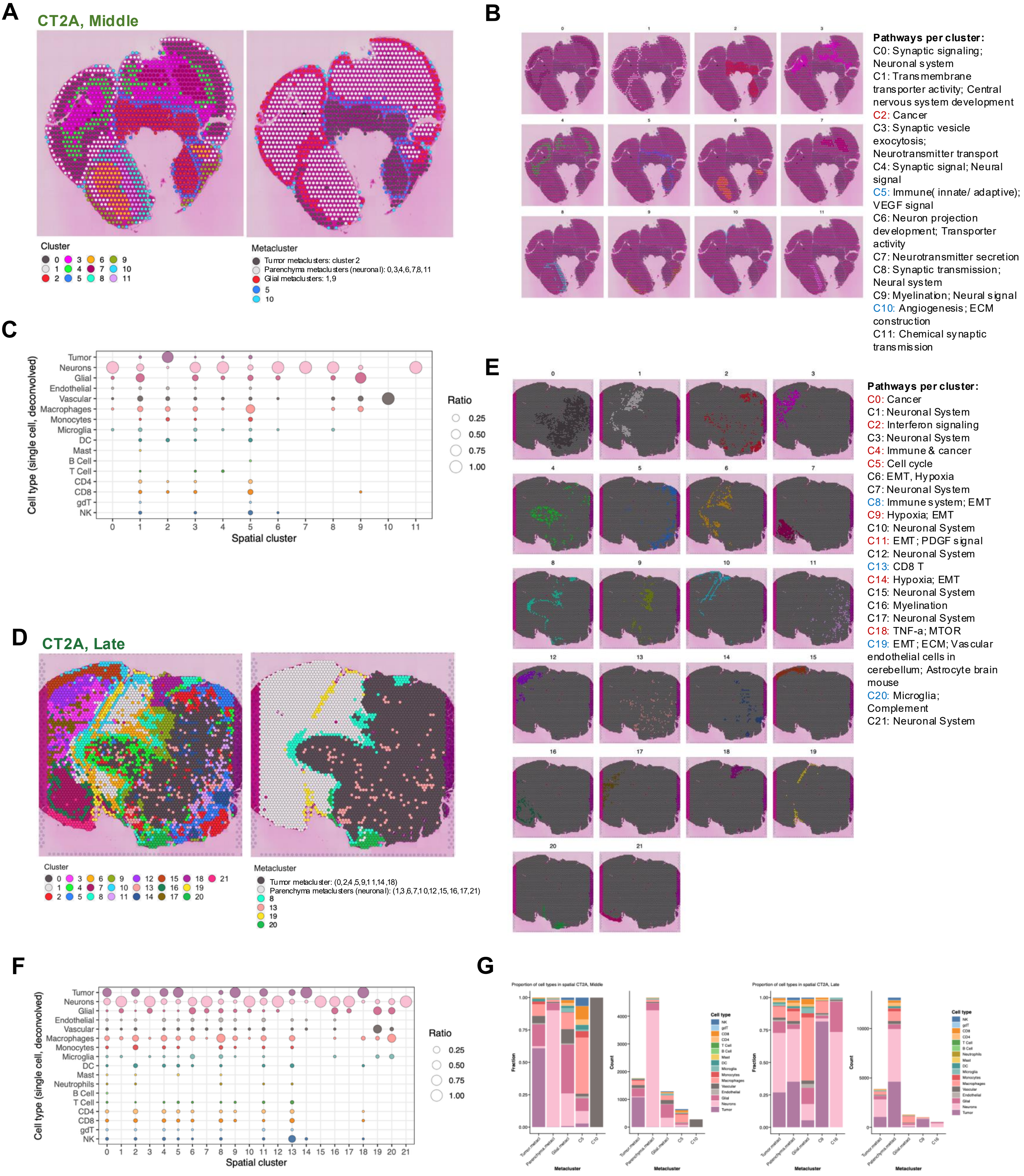
A
CT2A, Middle
Cluster
0 1 2 3 4 5 6 7 8 9 10 11
Metacluster
Tumor metaclusters: cluster 2
Parenchyma metaclusters (neuronal): 0,3,4,6,7,8,11
Glial metaclusters: 1,9
5
10
B
Pathways per cluster:
C0: Synaptic signaling; Neuronal system
C1: Transmembrane transporter activity; Central nervous system development
C2: Cancer
C3: Synaptic vesicle exocytosis; Neurotransmitter transport
C4: Synaptic signal; Neural signal
C5: Immune( innate/ adaptive); VEGF signal
C6: Neuron projection development; Transporter activity
C7: Neurotransmitter secretion
C8: Synaptic transmission; Neural system
C9: Myelination; Neural signal
C10: Angiogenesis; ECM construction
C11: Chemical synaptic transmission
C
Cell type (single cell, deconvolved)
Tumor
Neurons
Glial
Endothelial
Vascular
Macrophages
Monocytes
Microglia
DC
Mast
B Cell
T Cell
CD4
CD8
gdT
NK
Spatial cluster
Ratio
0.25
0.50
0.75
1.00
D
CT2A, Late
Cluster
0 1 2 3 4 5 6 7 8 9 10 11 12 13 14 15 16 17 18 19 20 21
Metacluster
Tumor metacluster: (0,2,4,5,9,11,14,18)
Parenchyma metaclusters (neuronal): (1,3,6,7,10,12,15,16,17,21)
8
13
19
20
E
Pathways per cluster:
C0: Cancer
C1: Neuronal System
C2: Interferon signaling
C3: Neuronal System
C4: Immune & cancer
C5: Cell cycle
C6: EMT, Hypoxia
C7: Neuronal System
C8: Immune system; EMT
C9: Hypoxia; EMT
C10: Neuronal System
C11: EMT; PDGF signal
C12: Neuronal System
C13: CD8 T
C14: Hypoxia; EMT
C15: Neuronal System
C16: Myelination
C17: Neuronal System
C18: TNF-a; MTOR
C19: EMT; ECM; Vascular endothelial cells in cerebellum; Astrocyte brain mouse
C20: Microglia; Complement
C21: Neuronal System
F
Cell type (single cell, deconvolved)
Tumor
Neurons
Glial
Endothelial
Vascular
Macrophages
Monocytes
Microglia
DC
Mast
Neutrophils
B Cell
T Cell
CD4
CD8
gdT
NK
Spatial cluster
Ratio
0.25
0.50
0.75
1.00
G
Proportion of cell types in spatial CT2A, Middle
Proportion of cell types in spatial CT2A, Late
Fraction
Count
Metacluster
Cell type

**Supplementary Figure S3. Functional annotation and cell type composition of CT2A spatial compartments along tumor progression.**

**A, D.** Spatial distribution and cluster composition of CT2A (A) middle and (D) late-stage samples. **B, E.** Functional annotation of spatial clusters for (B) middle and (E) late-stage samples. Annotation was performed via over-representation analysis of cluster-specific markers; tumor-associated clusters are indicated in red, neuro-glial clusters in black, and immune or vascular clusters in blue. **C, F.** Dot plots showing the proportions of scRNA-seq-defined cell types projected onto spatial clusters for (C) middle and (F) late-stage samples following CellTrek deconvolution.

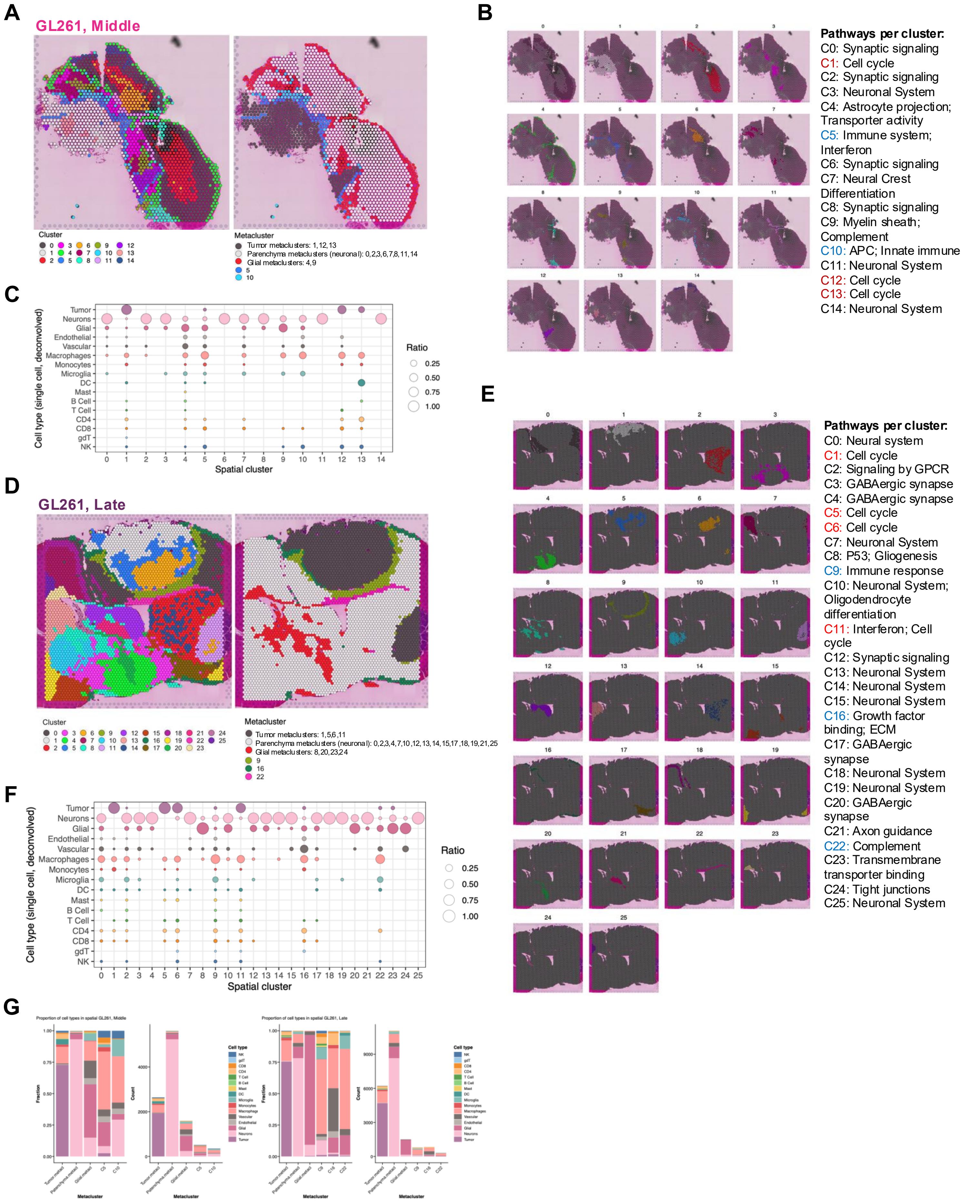
A
GL261, Middle
Cluster
0 1 2 3 4 5 6 7 8 9 10 11 12 13 14
Metacluster
Tumor metaclusters: 1,12,13
Parenchyma metaclusters (neuronal): 0,2,3,6,7,8,11,14
Glial metaclusters: 4,9
5
10
B
0 1 2 3 4 5 6 7 8 9 10 11 12 13 14
Pathways per cluster:
C0: Synaptic signaling
C1: Cell cycle
C2: Synaptic signaling
C3: Neuronal System
C4: Astrocyte projection; Transporter activity
C5: Immune system; Interferon
C6: Synaptic signaling
C7: Neural Crest Differentiation
C8: Synaptic signaling
C9: Myelin sheath; Complement
C10: APC; Innate immune
C11: Neuronal System
C12: Cell cycle
C13: Cell cycle
C14: Neuronal System
C
Tumor
Neurons
Glial
Endothelial
Vascular
Macrophages
Monocytes
Microglia
DC
Mast
B Cell
T Cell
CD4
CD8
gdT
NK
Cell type (single cell, deconvolved)
0 1 2 3 4 5 6 7 8 9 10 11 12 13 14
Spatial cluster
Ratio
0.25
0.50
0.75
1.00
D
GL261, Late
Cluster
0 1 2 3 4 5 6 7 8 9 10 11 12 13 14 15 16 17 18 19 20 21 22 23 24 25
Metacluster
Tumor metaclusters: 1,5,6,11
Parenchyma metaclusters (neuronal): 0,2,3,4,7,10,12,13,14,15,17,18,19,21,25
Glial metaclusters: 8,20,23,24
9
16
22
E
0 1 2 3 4 5 6 7 8 9 10 11 12 13 14 15 16 17 18 19 20 21 22 23 24 25
Pathways per cluster:
C0: Neural system
C1: Cell cycle
C2: Signaling by GPCR
C3: GABAergic synapse
C4: GABAergic synapse
C5: Cell cycle
C6: Cell cycle
C7: Neuronal System
C8: P53; Gliogenesis
C9: Immune response
C10: Neuronal System; Oligodendrocyte differentiation
C11: Interferon; Cell cycle
C12: Synaptic signaling
C13: Neuronal System
C14: Neuronal System
C15: Neuronal System
C16: Growth factor binding; ECM
C17: GABAergic synapse
C18: Neuronal System
C19: Neuronal System
C20: GABAergic synapse
C21: Axon guidance
C22: Complement
C23: Transmembrane transporter binding
C24: Tight junctions
C25: Neuronal System
F
Tumor
Neurons
Glial
Endothelial
Vascular
Macrophages
Monocytes
Microglia
DC
Mast
B Cell
T Cell
CD4
CD8
gdT
NK
Cell type (single cell, deconvolved)
0 1 2 3 4 5 6 7 8 9 10 11 12 13 14 15 16 17 18 19 20 21 22 23 24 25
Spatial cluster
Ratio
0.25
0.50
0.75
1.00
G
Proportion of cell types in spatial GL261, Middle
Proportion of cell types in spatial GL261, Late
Fraction
Count
Metacluster
Tumor.metacl
Parenchyma.metacl
Glial.metacl
C5
C10
C9
C16
C22
Cell type
NK
gdT
CD8
CD4
T Cell
B Cell
Mast
DC
Microglia
Monocytes
Macrophages
Vascular
Endothelial
Glial
Neurons
Tumor

**Supplementary Figure S4. Functional annotation and cell type composition of GL261 spatial compartments along tumor progression.**

**A, D.** Spatial distribution and cluster composition of GL261 (A) middle and (D) late-stage samples. **B, E.** Functional annotation of spatial clusters for (B) middle and (E) late-stage samples. Annotation was performed via over-representation analysis of cluster-specific markers; tumor-associated clusters are indicated in red, neuro-glial clusters in black, and immune or vascular clusters in blue. **C, F.** Dot plots showing the proportions of scRNA-seq-defined cell types projected onto spatial clusters for (C) middle and (F) late-stage samples following CellTrek deconvolution.

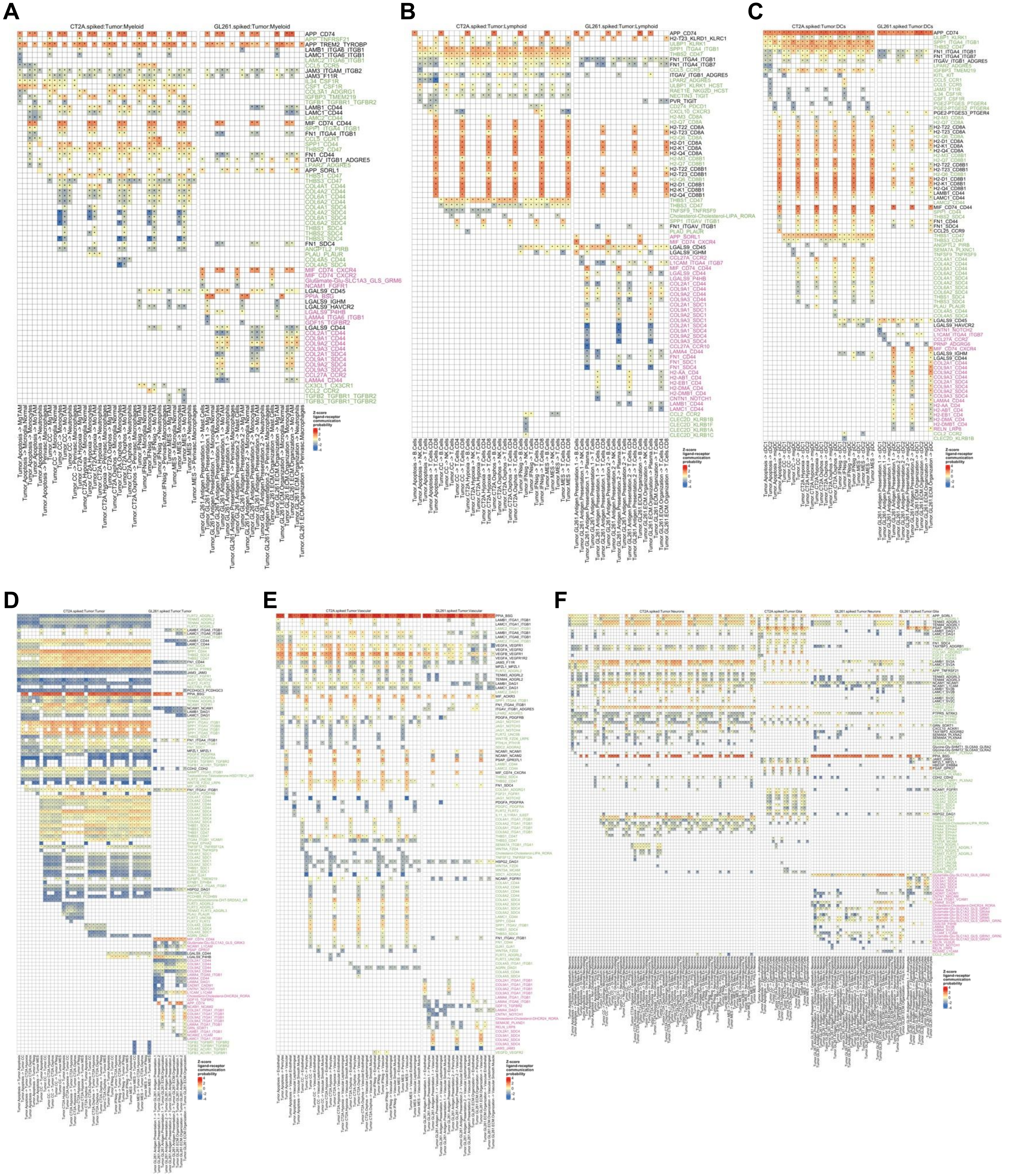
A
CT2A.spiked:Tumor:Myeloid
GL261.spiked:Tumor:Myeloid
B
CT2A.spiked:Tumor:Lymphoid
GL261.spiked:Tumor:Lymphoid
C
CT2A.spiked:Tumor:DCs
GL261.spiked:Tumor:DCs
D
CT2A.spiked:Tumor:Tumor
GL261.spiked:Tumor:Tumor
E
CT2A.spiked:Tumor:Vascular
GL261.spiked:Tumor:Vascular
F
CT2A.spiked:Tumor:Neurons
CT2A.spiked:Tumor:Glia
GL261.spiked:Tumor:Neurons
GL261.spiked:Tumor:Glia
Z-score ligand-receptor communication probability

**Supplementary Figure S5. Model-specific and shared ligand-receptor interactions between tumor cells and the microenvironment.**

Heatmaps display z-scored interaction probabilities for significant ligand-receptor (L-R) pairs, with tumor cells as "sender" and specific cell populations as "recipients." Probabilities were calculated per model using CellChat; z-scores are relative to each model's total significant interactions. L-R labels indicate interactions unique to CT2A (green), unique to GL261 (pink), or shared by both models (black). Subsets represent interactions between tumor cells and: **(A)** myeloid cells, **(B)** lymphoid cells, **(C)** dendritic cells, **(D)** tumor cells (autocrine), **(E)** vasculature-like cells, and **(F)** neural and glial cells.

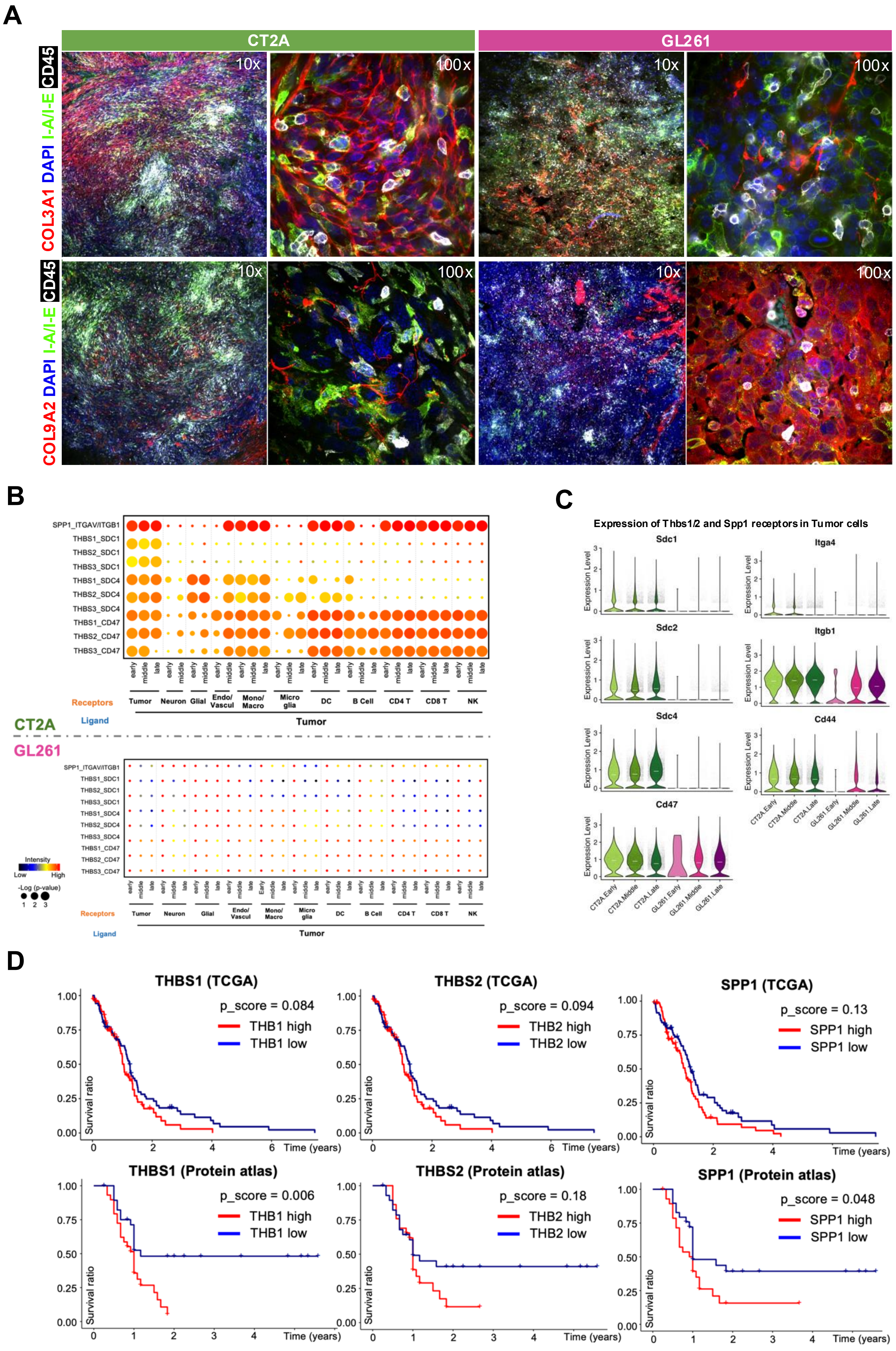
A
CT2A
GL261
CD45 I-A/I-E DAPI COL3A1
CD45 I-A/I-E DAPI COL9A2
10x
100x
B
CT2A
GL261
Receptors
Ligand
Tumor
Neuron
Glial
Endo/ Vascul
Mono/ Macro
Micro glia
DC
B Cell
CD4 T
CD8 T
NK
SPP1_ITGAV/ITGB1
THBS1_SDC1
THBS2_SDC1
THBS3_SDC1
THBS1_SDC4
THBS2_SDC4
THBS3_SDC4
THBS1_CD47
THBS2_CD47
THBS3_CD47
Intensity
Low
High
-Log (p-value)
C
Expression of Thbs1/2 and Spp1 receptors in Tumor cells
Sdc1
Itga4
Sdc2
Itgb1
Sdc4
Cd44
Cd47
Expression Level
CT2A.Early
CT2A.Middle
CT2A.Late
GL261.Early
GL261.Middle
GL261.Late
D
THBS1 (TCGA)
p_score = 0.084
THB1 high
THB1 low
THBS2 (TCGA)
p_score = 0.094
THB2 high
THB2 low
SPP1 (TCGA)
p_score = 0.13
SPP1 high
SPP1 low
THBS1 (Protein atlas)
p_score = 0.006
THB1 high
THB1 low
THBS2 (Protein atlas)
p_score = 0.18
THB2 high
THB2 low
SPP1 (Protein atlas)
p_score = 0.048
SPP1 high
SPP1 low
Survival ratio
Time (years)

**Supplementary Figure S6. Validation of divergent extracellular matrix programs and clinical correlations with the Thbs/Spp1 axis.**

**A.** Representative immunofluorescence (IF) micrographs of late-stage orthotopic CT2A (left) and GL261 (right) tumors. The top row displays COL3A1 deposition (red) alongside I-A/I-E (MHC-II, green), CD45 (white), and DAPI nuclei counterstain (blue). The bottom row displays COL9A2 deposition (red) alongside the same markers. Corresponding 10X overview and 100X detailed views are presented, visualizing differential patterns of extracellular matrix architecture and MHC-II expression between the two models. **B.** Dot plots representing interaction intensity (color) and significance (-log(p-value)) of tumor-to-microenvironment ligand-receptor pairs in CT2A and GL261 across sampled timepoints. **C.** Expression of receptors for THBS1/2 and SPP1 in tumoe cells along time. **D.** Kaplan–Meier survival analysis of glioblastoma patients stratified by high (above median) or low (below median) expression of THBS1, THBS2, and SPP1. Data are presented for TCGA (top) and The Human Protein Atlas (bottom) cohorts.

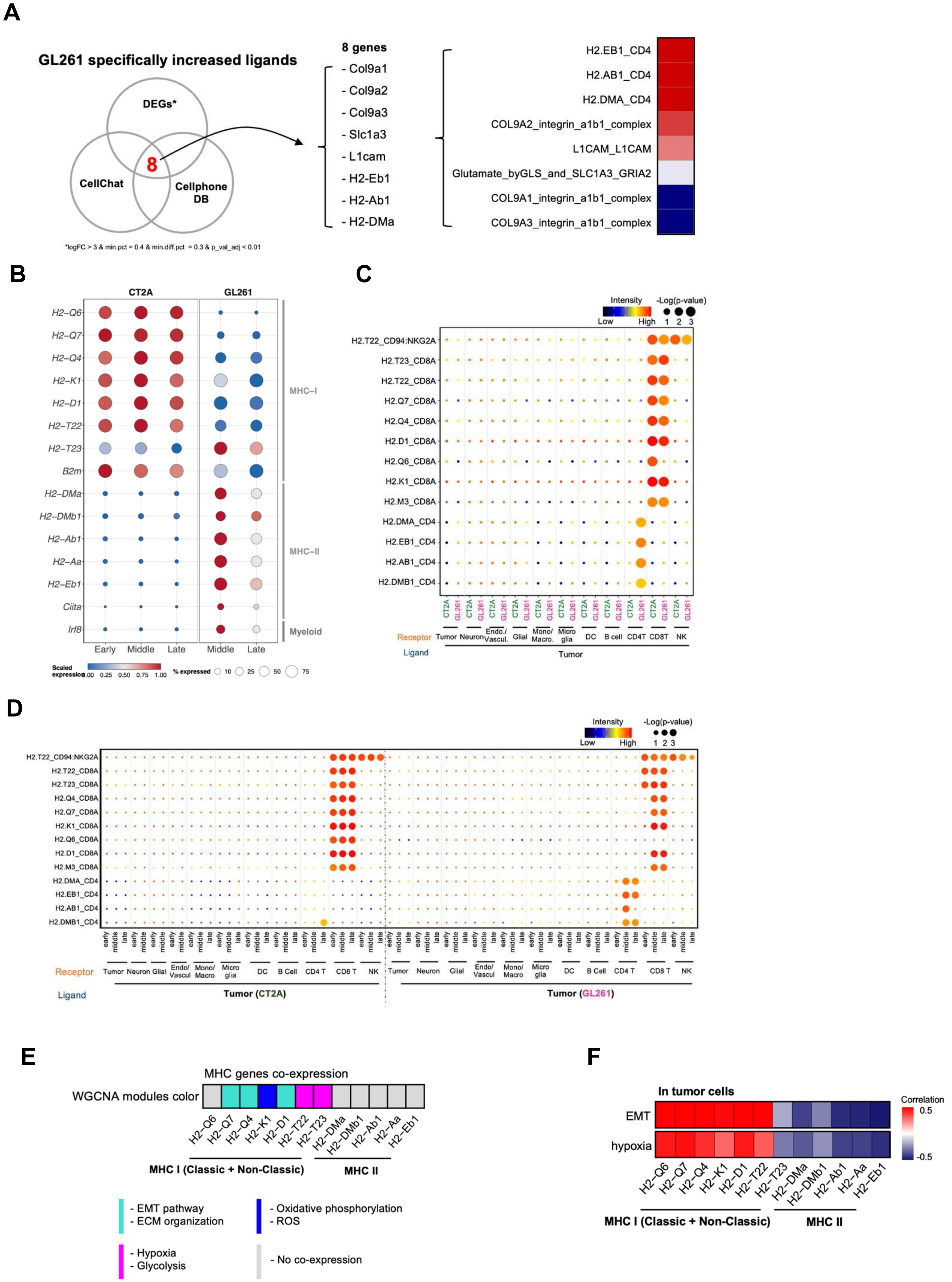
A
GL261 specifically increased ligands
DEGs*
CellChat
Cellphone DB
8
*logFC > 3 & min.pct = 0.4 & min.diff.pct = 0.3 & p_val_adj < 0.01
8 genes
- Col9a1
- Col9a2
- Col9a3
- Slc1a3
- L1cam
- H2-Eb1
- H2-Ab1
- H2-DMa
H2.EB1_CD4
H2.AB1_CD4
H2.DMA_CD4
COL9A2_integrin_a1b1_complex
L1CAM_L1CAM
Glutamate_byGLS_and_SLC1A3_GRIA2
COL9A1_integrin_a1b1_complex
COL9A3_integrin_a1b1_complex
B
CT2A
GL261
H2–Q6
H2–Q7
H2–Q4
H2–K1
H2–D1
H2–T22
H2–T23
B2m
H2–DMa
H2–DMb1
H2–Ab1
H2–Aa
H2–Eb1
Ciita
Irf8
MHC–I
MHC–II
Myeloid
Early
Middle
Late
Middle
Late
Scaled expression
0.00 0.25 0.50 0.75 1.00
% expressed
10 25 50 75
C
Intensity
Low
High
-Log(p-value)
1 2 3
H2.T22_CD94:NKG2A
H2.T23_CD8A
H2.T22_CD8A
H2.Q7_CD8A
H2.Q4_CD8A
H2.D1_CD8A
H2.Q6_CD8A
H2.K1_CD8A
H2.M3_CD8A
H2.DMA_CD4
H2.EB1_CD4
H2.AB1_CD4
H2.DMB1_CD4
CT2A
GL261
Receptor
Ligand
Tumor
Neuron
Endo./ Vascul.
Glial
Mono/ Macro.
Micro glia
DC
B cell
CD4T
CD8T
NK
Tumor
D
Intensity
Low
High
-Log(p-value)
1 2 3
H2.T22_CD94:NKG2A
H2.T22_CD8A
H2.T23_CD8A
H2.Q4_CD8A
H2.Q7_CD8A
H2.K1_CD8A
H2.Q6_CD8A
H2.D1_CD8A
H2.M3_CD8A
H2.DMA_CD4
H2.EB1_CD4
H2.AB1_CD4
H2.DMB1_CD4
early
middle
late
Receptor
Tumor
Neuron
Glial
Endo/ Vascul
Mono/ Macro
Micro glia
DC
B Cell
CD4 T
CD8 T
NK
Ligand
Tumor (CT2A)
Tumor (GL261)
E
MHC genes co-expression
WGCNA modules color
H2-Q6
H2-Q7
H2-Q4
H2-K1
H2-D1
H2-T22
H2-T23
H2-DMa
H2-DMb1
H2-Ab1
H2-Aa
H2-Eb1
MHC I (Classic + Non-Classic)
MHC II
- EMT pathway
- ECM organization
- Oxidative phosphorylation
- ROS
- Hypoxia
- Glycolysis
- No co-expression
F
In tumor cells
EMT
hypoxia
Correlation
0.5
-0.5
H2-Q6
H2-Q7
H2-Q4
H2-K1
H2-D1
H2-T22
H2-T23
H2-DMa
H2-DMb1
H2-Ab1
H2-Aa
H2-Eb1
MHC I (Classic + Non-Classic)
MHC II

**Supplementary Figure S7. Characterization of MHC-mediated signaling.**

**A.** Venn diagram (left) identifying GL261-enriched ligands consistently detected by both CellChat and CellPhoneDB; heatmap (right) illustrates the relative interaction intensities for these ligand-receptor pairs**. B.** Dot plot illustrating H2 (MHC Class I and II) gene expression across tumor cell populations. **C, D.** Tumor-to-microenvironment ligand-receptor (L-R) interactions utilizing MHC molecules as ligands. Comparisons are presented for (C) bulk models and (D) timepoints. **E.** Gene co-expression modules and enriched biological pathways associated with MHC genes, as identified by weighted gene co-expression network analysis (WGCNA). **F.** Heatmap representing correlation scores between MHC gene expression and biological pathway scores for EMT (top) and hypoxia (bottom

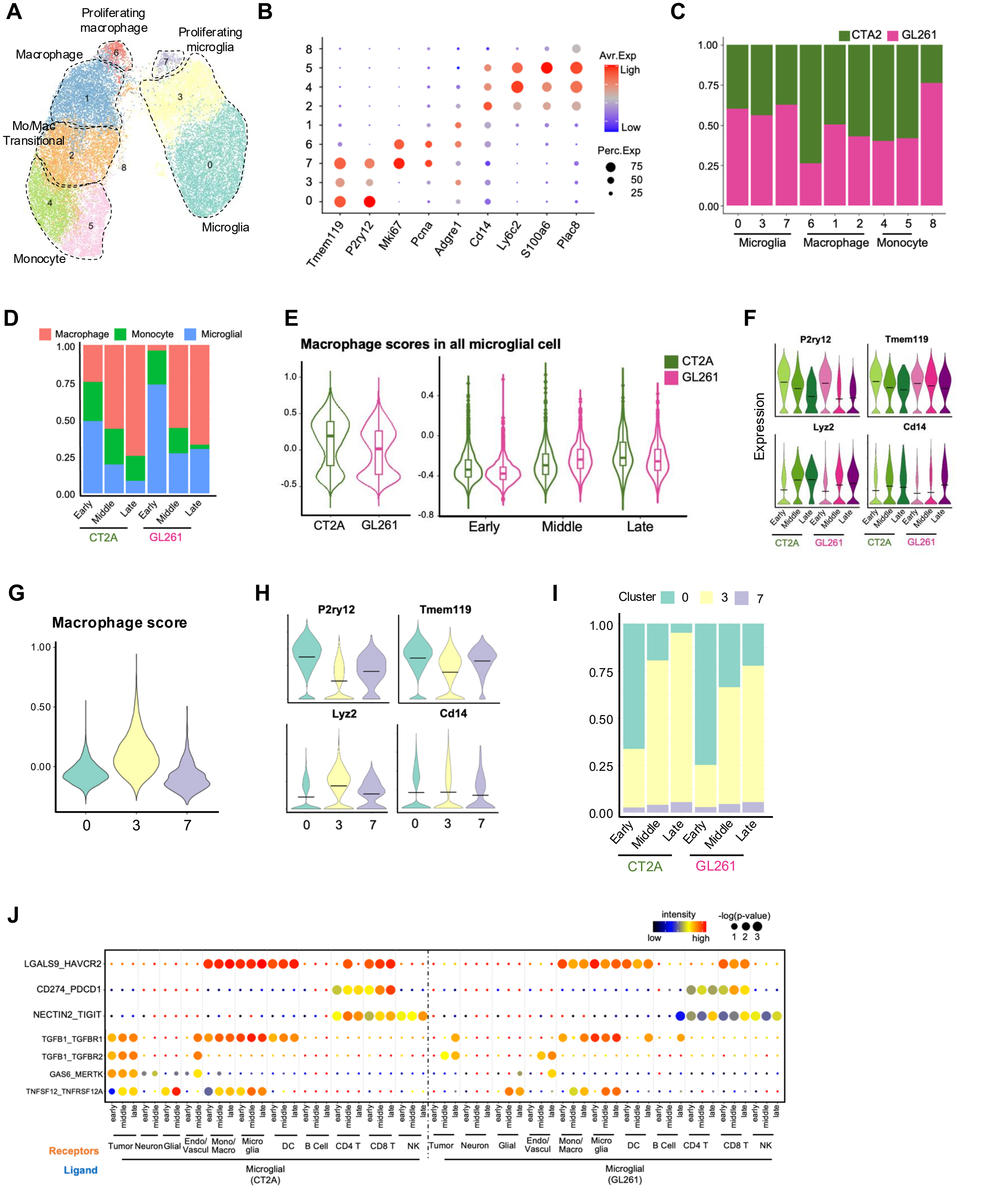

A
Proliferating macrophage
Proliferating microglia
Macrophage
Mo/Mac Transitional
Microglia
Monocyte
B
Avr.Exp
Ligh
Low
Perc.Exp
75
50
25
Tmem119
P2ry12
Mki67
Pcna
Adgre1
Cd14
Ly6c2
S100a6
Plac8
C
CTA2
GL261
Microglia
Macrophage
Monocyte
D
Macrophage
Monocyte
Microglial
Early
Middle
Late
CT2A
GL261
E
Macrophage scores in all microglial cell
CT2A
GL261
Early
Middle
Late
F
P2ry12
Tmem119
Lyz2
Cd14
Expression
G
Macrophage score
H
P2ry12
Tmem119
Lyz2
Cd14
I
Cluster
J
intensity
low
high
-log(p-value)
LGALS9_HAVCR2
CD274_PDCD1
NECTIN2_TIGIT
TGFB1_TGFBR1
TGFB1_TGFBR2
GAS6_MERTK
TNFSF12_TNFRSF12A
early
middle
late
Tumor
Neuron
Glial
Endo/ Vascul
Mono/ Macro
Micro glia
DC
B Cell
CD4 T
CD8 T
NK
Receptors
Ligand
Microglial (CT2A)
Microglial (GL261)

**Supplementary Figure S8. Microglial cells and macrophages evolution in two tumor models over time.**

**A.** Sub-clustering of the myeloid compartment. UMAP projection illustrating high-resolution clustering of macrophages, monocytes, and microglia. **B.** Expression of canonical markers defining myeloid cell identity. **C, D.** Proportional distribution of myeloid cell types. Comparisons are presented between (C) tumor models and (D) across the global myeloid subset. **E.** Quantification of macrophage-state signature scores. Scores are based on macrophage-specific gene expression relative to monocytes and microglia, compared between models (left) and across timepoints (right). **F.** Temporal expression profiles of microglial (top) and macrophage (bottom) markers across tumor progression. **G, H.** Evaluation of macrophage-like features within microglial sub-clusters. **(G)** Macrophage-state signature scores and **(H)** canonical marker expression across three identified microglial subtypes. **I.** Proportional dynamics of microglial sub-clusters across tumor models and timepoints. **J.** Microglia-mediated ligand-receptor interactions. Dot size and color represent $-\log_{10}$(p-value) and interaction intensity, respectively, between microglial ligands and recipient cell receptors.

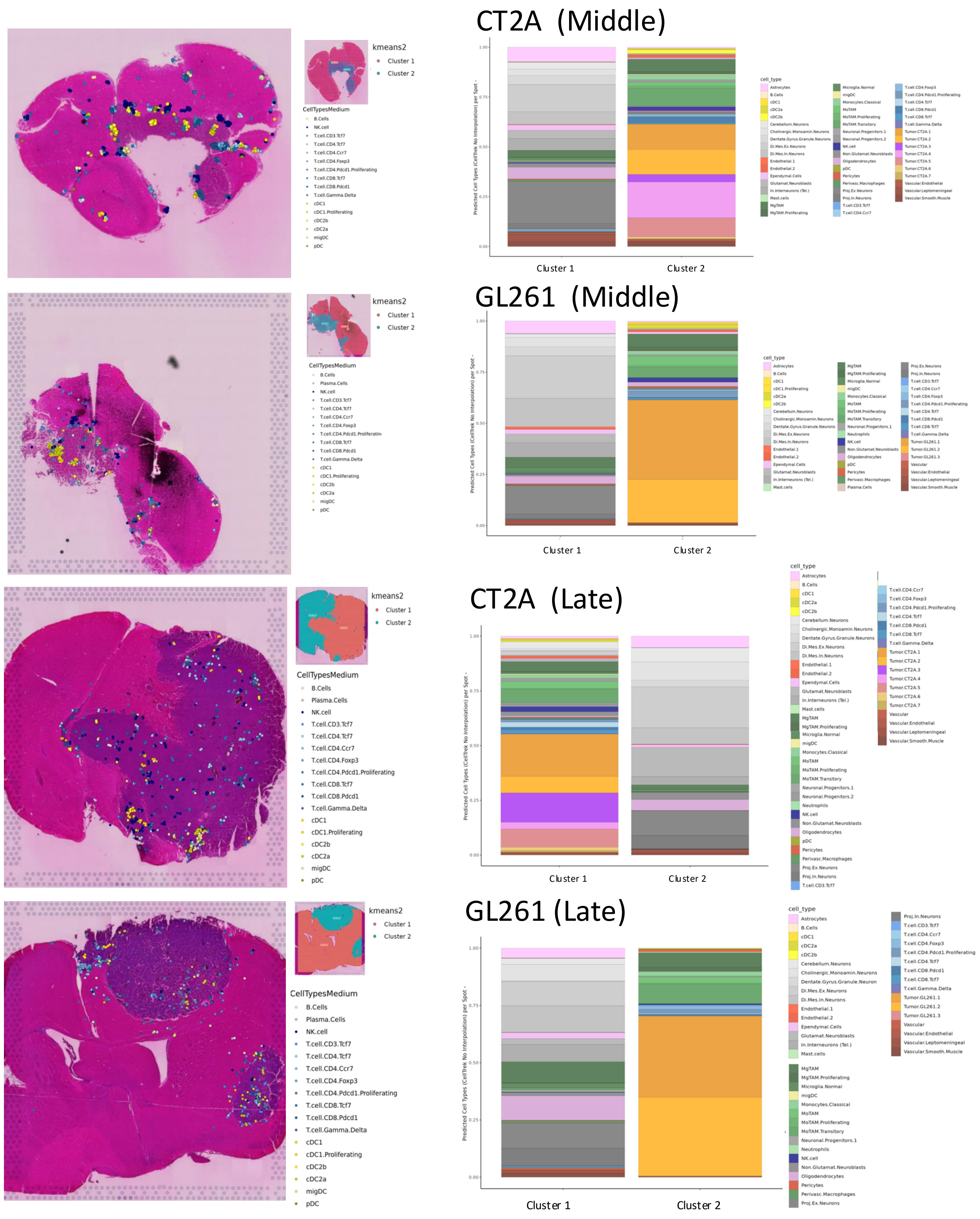

CT2A (Middle)
GL261 (Middle)
CT2A (Late)
GL261 (Late)
kmeans2
Cluster 1
Cluster 2
CellTypesMedium
B.Cells
Plasma.Cells
NK.cell
T.cell.CD3.Tcf7
T.cell.CD4.Tcf7
T.cell.CD4.Ccr7
T.cell.CD4.Foxp3
T.cell.CD4.Pdcd1.Proliferating
T.cell.CD8.Tcf7
T.cell.CD8.Pdcd1
T.cell.Gamma.Delta
cDC1
cDC1.Proliferating
cDC2b
cDC2a
migDC
pDC
Predicted Cell Types (CellTrek No Interpolation) per Spot
cell_type
Astrocytes
B.Cells
cDC1
cDC2a
cDC2b
Cerebellum.Neurons
Cholinergic.Monoamin.Neurons
Dentate.Gyrus.Granule.Neurons
Di.Mes.Ex.Neurons
Di.Mes.In.Neurons
Endothelial.1
Endothelial.2
Ependymal.Cells
Glutamat.Neuroblasts
In.Interneurons (Tel.)
Mast.cells
MgTAM
MgTAM.Proliferating
Microglia.Normal
migDC
Monocytes.Classical
MoTAM
MoTAM.Proliferating
MoTAM.Transitory
Neuronal.Progenitors.1
Neuronal.Progenitors.2
Neutrophils
NK.cell
Non.Glutamat.Neuroblasts
Oligodendrocytes
pDC
Pericytes
Perivasc.Macrophages
Proj.Ex.Neurons
Proj.In.Neurons
Tumor.CT2A.1
Tumor.CT2A.2
Tumor.CT2A.3
Tumor.CT2A.4
Tumor.CT2A.5
Tumor.CT2A.6
Tumor.CT2A.7
Tumor.GL261.1
Tumor.GL261.2
Tumor.GL261.3
Vascular
Vascular.Endothelial
Vascular.Leptomeningeal
Vascular.Smooth.Muscle
0.00
0.25
0.50
0.75
1.00

**Supplementary Figure S9. Proportions of high-resolution cell populations in tumor and parenchyma clusters, as determined by spatial ranger K-means clustering.**

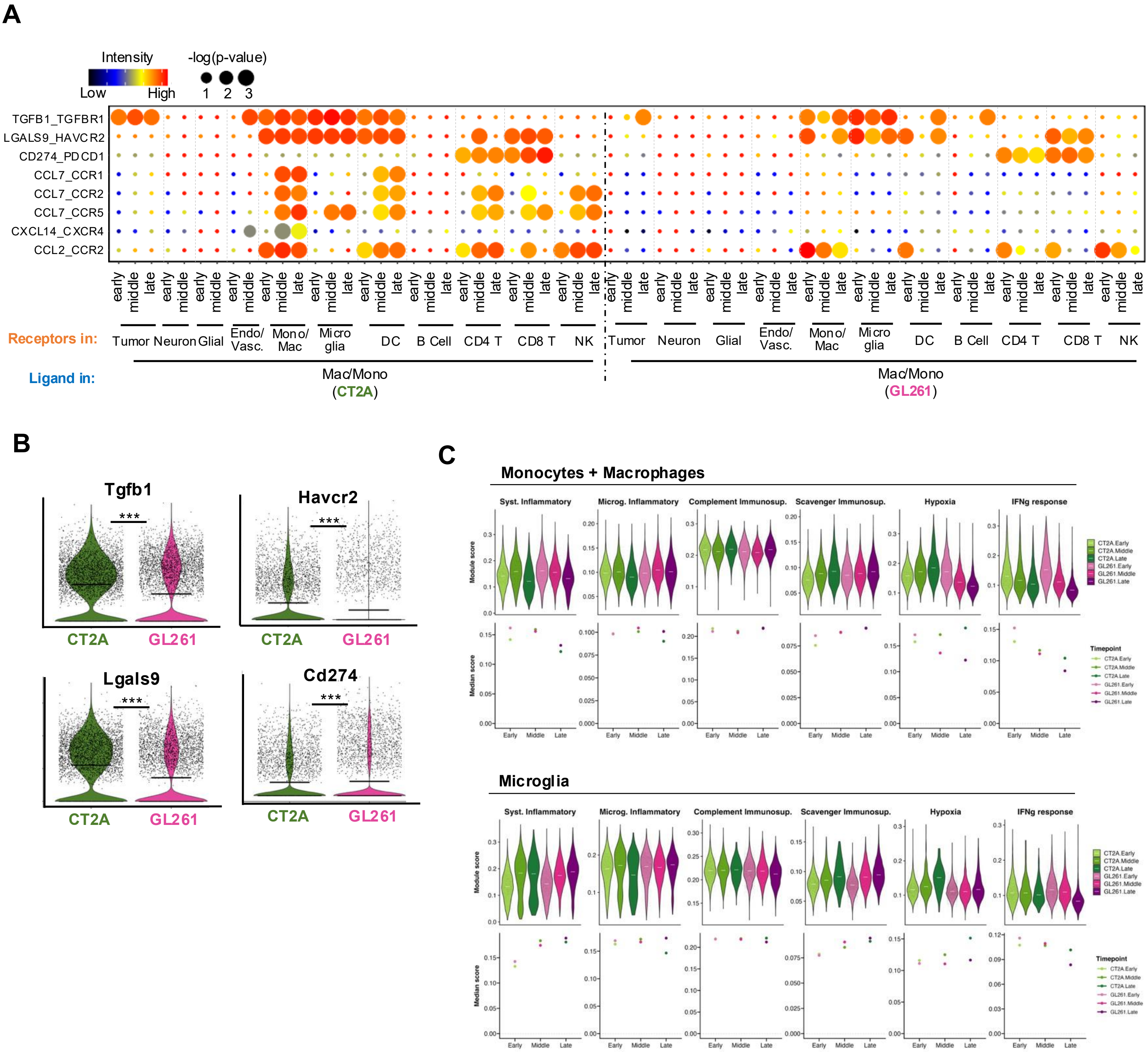
A
Intensity
Low
High
-log(p-value)
1 2 3
TGFB1_TGFBR1
LGALS9_HAVCR2
CD274_PDCD1
CCL7_CCR1
CCL7_CCR2
CCL7_CCR5
CXCL14_CXCR4
CCL2_CCR2
early middle late
Receptors in:
Tumor Neuron Glial Endo/ Vasc. Mono/ Mac Micro glia DC B Cell CD4 T CD8 T NK
Ligand in:
Mac/Mono (CT2A)
Mac/Mono (GL261)
B
Tgfb1
Havcr2
Lgals9
Cd274
***
CT2A
GL261
C
Monocytes + Macrophages
Microglia
Syst. Inflammatory
Microg. Inflammatory
Complement Immunosup.
Scavenger Immunosup.
Hypoxia
IFNg response
Module score
Median score
Early
Middle
Late
Timepoint
CT2A.Early
CT2A.Middle
CT2A.Late
GL261.Early
GL261.Middle
GL261.Late

**Supplementary Figure S10. Spatiotemporal dynamics of macrophage-mediated signaling and immunomodulatory profiles.**

**A.** Macrophage-to-microenvironment ligand-receptor interactions across tumor progression. Dot size and color represent $-\log_{10}$(p-value) and interaction intensity, respectively, for macrophage-derived ligands and their corresponding receptors in CT2A and GL261. **B.** Expression of immunomodulatory and checkpoint genes. Violin plots illustrate expression levels of *Tgfb1*, *Havcr2*, *Lgals9*, and *Cd274* in macrophages between tumor models. **C.** Distribution of macrophage immunomodulatory metaprograms (Miller *et al.*, 2015) scores across monocyte + macrophages (top) and microglia (bottom) over time.

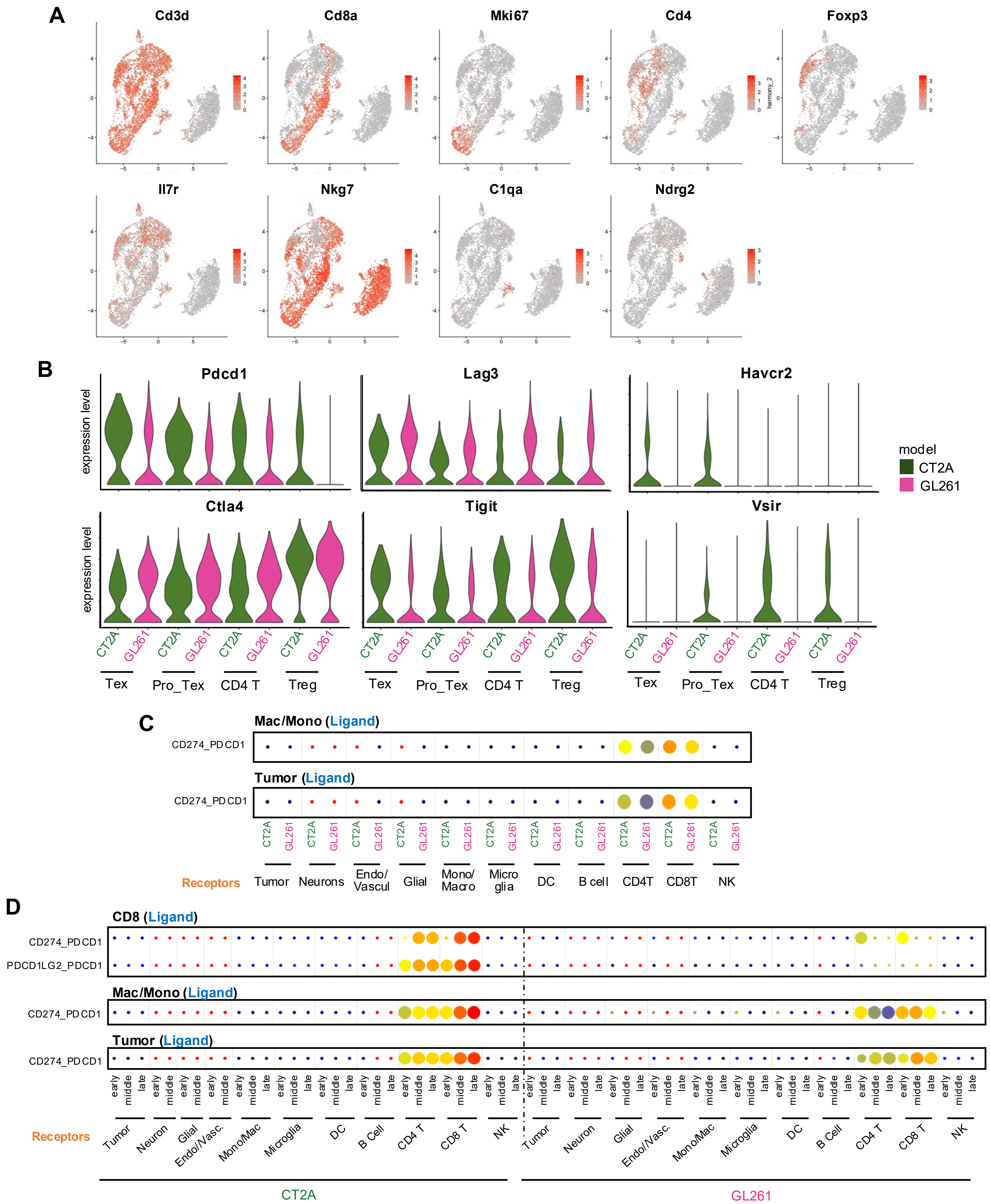

A
Cd3d
Cd8a
Mki67
Cd4
Foxp3
Il7r
Nkg7
C1qa
Ndrg2
B
Pdcd1
Lag3
Havcr2
Ctla4
Tigit
Vsir
expression level
model
CT2A
GL261
Tex
Pro_Tex
CD4 T
Treg
C
Mac/Mono (Ligand)
Tumor (Ligand)
CD274_PDCD1
Receptors
Tumor
Neurons
Endo/ Vascul
Glial
Mono/ Macro
Micro glia
DC
B cell
CD4T
CD8T
NK
D
CD8 (Ligand)
Mac/Mono (Ligand)
Tumor (Ligand)
CD274_PDCD1
PDCD1LG2_PDCD1
early
middle
late
Receptors
Tumor
Neuron
Glial
Endo/Vasc.
Mono/Mac
Microglia
DC
B Cell
CD4 T
CD8 T
NK
CT2A
GL261

**Supplementary Figure S11. Dysfunctional activity of CD8 cells between CT2A and GL261.**

**A.** Feature plot indicating expression levels of marker genes corresponding to each cell type. **B.** Violin plot showing expression levels of dysfunction markers genes in CD4 and CD8 T cells between CT2A and GL261. **C.** CD274_PDCD1 interaction between macrophage and T lymphocytes (top) and between cancer cells and T lymphocytes (bottom) in two tumor models. Dot size and color represent $-\log_{10}$(p-value) and interaction intensity, respectively. **D.** CD274_PDCD1 and PDCD1LG2_PDCD1 interaction between immune-immune and immune-cancer cells across time points. Dot size and color represent $-\log_{10}$(p-value) and interaction intensity, respectively.

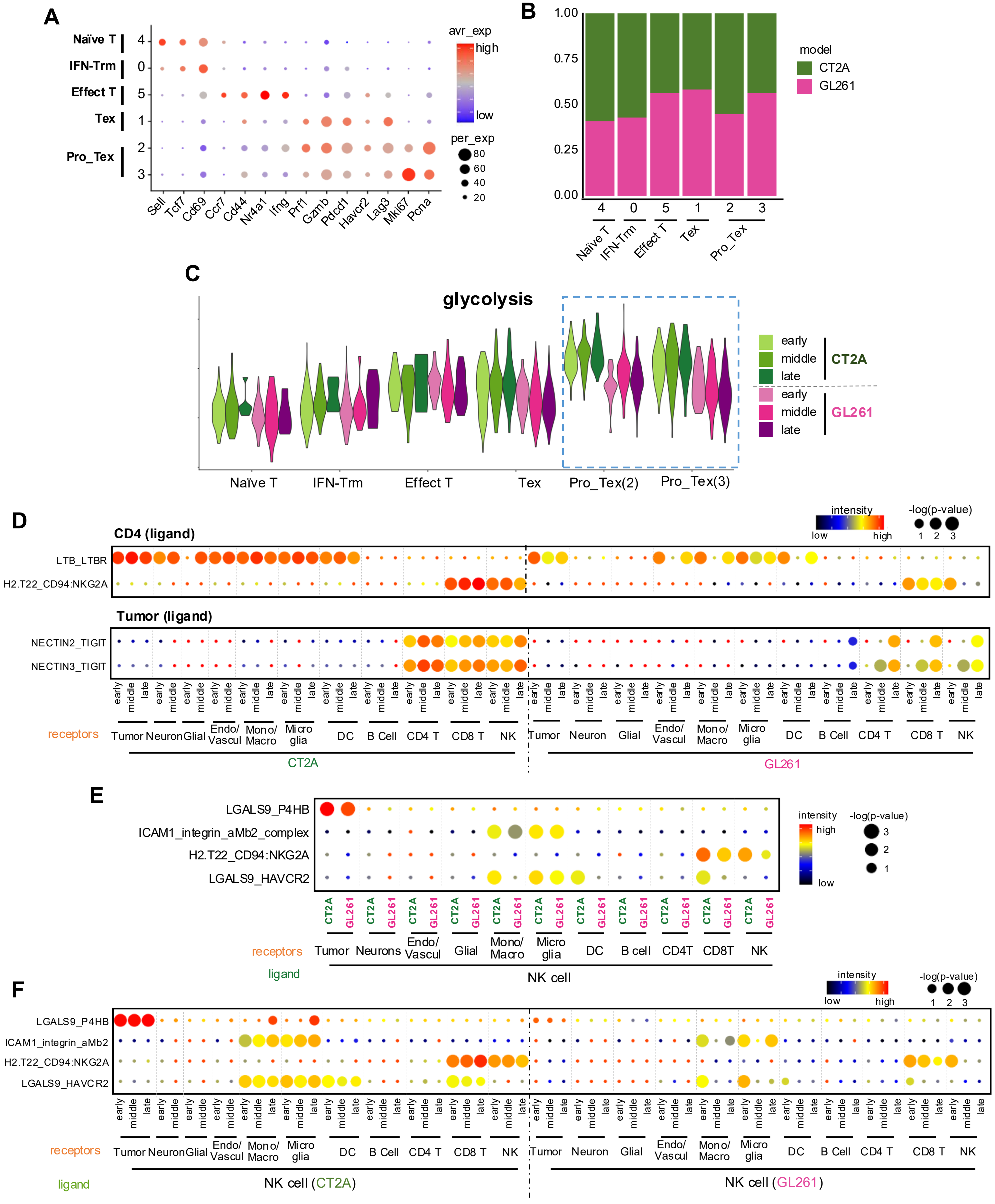
A
Naïve T
IFN-Trm
Effect T
Tex
Pro_Tex
4
0
5
1
2
3
Sell
Tcf7
Cd69
Ccr7
Cd44
Nr4a1
Ifng
Prf1
Gzmb
Pdcd1
Havcr2
Lag3
Mki67
Pcna
avr_exp
high
low
per_exp
80
60
40
20
B
1.00
0.75
0.50
0.25
0.00
model
CT2A
GL261
C
glycolysis
early
middle
late
CT2A
early
middle
late
GL261
Naïve T
IFN-Trm
Effect T
Tex
Pro_Tex(2)
Pro_Tex(3)
D
CD4 (ligand)
LTB_LTBR
H2.T22_CD94:NKG2A
Tumor (ligand)
NECTIN2_TIGIT
NECTIN3_TIGIT
intensity
low
high
-log(p-value)
1
2
3
receptors
Tumor
Neuron
Glial
Endo/ Vascul
Mono/ Macro
Micro glia
DC
B Cell
CD4 T
CD8 T
NK
CT2A
GL261
E
LGALS9_P4HB
ICAM1_integrin_aMb2_complex
H2.T22_CD94:NKG2A
LGALS9_HAVCR2
Neurons
Glial
B cell
CD4T
CD8T
ligand
NK cell
F
ICAM1_integrin_aMb2
NK cell (CT2A)
NK cell (GL261)

**Supplementary Figure S12. Immune evasive interaction by lymphocytes.**

**A.** Dop plot showing representative marker gene expression for each cell type. **B.** Bar plot showing cell population between two tumor models for each cell type. **C.** Violin plot displaying glycolysis scores between CT2A and GL261 across CD8 T cell subtypes.. **D.** Cell-cell interaction between CT2A and GL261, with CT4 cells (top) or cancer cells (bottom) acting as ligands. Dot size and color represent -log(p-value) and interaction intensity, respectively. **E,F.** Cell-cell interaction between CT2A and GL261(E) and across time pointes (F), with NK cells acting as ligands.
